# Single Cell RNA Sequencing Reveals Emergent Notochord-Derived Cell Subpopulations in the Postnatal Nucleus Pulposus

**DOI:** 10.1101/2023.05.21.541589

**Authors:** Chenghao Zhang, Leilei Zhong, Yian Khai Lau, Meilun Wu, Lutian Yao, Thomas P. Schaer, Robert L. Mauck, Neil R. Malhotra, Ling Qin, Lachlan J. Smith

## Abstract

Intervertebral disc degeneration is a leading cause of chronic low back pain. Cell-based strategies that seek to treat disc degeneration by regenerating the central nucleus pulposus hold significant promise, but key challenges remain. One of these is the inability of therapeutic cells to effectively mimic the performance of native nucleus pulposus cells, which are unique amongst skeletal cell types in that they arise from the embryonic notochord. In this study we use single cell RNA sequencing to demonstrate emergent heterogeneity amongst notochord-derived nucleus pulposus cells in the postnatal mouse disc. Specifically, we established the existence of early and late stage nucleus pulposus cells, corresponding to notochordal progenitor and mature cells, respectively. Late stage cells exhibited significantly higher expression levels of extracellular matrix genes including aggrecan, and collagens II and VI, along with elevated TGF-β and PI3K-Akt signaling. Additionally, we identified Cd9 as a novel surface marker of late stage nucleus pulposus cells, and demonstrated that these cells were localized to the nucleus pulposus periphery, increased in numbers with increasing postnatal age, and co-localized with emerging glycosaminoglycan-rich matrix. Finally, we used a goat model to show the Cd9+ nucleus pulposus cell numbers decrease with moderate severity disc degeneration, suggesting that these cells are associated with maintenance of the healthy nucleus pulposus extracellular matrix. Improved understanding of the developmental mechanisms underlying regulation of ECM deposition in the postnatal NP may inform improved regenerative strategies for disc degeneration and associated low back pain.

## Introduction

Lumbar intervertebral disc degeneration is strongly implicated as a cause of low back pain, the leading cause of disability worldwide [1–4]. The discs are the partially-movable joints of the spine, and each is comprised of three main substructures: a central, proteoglycan-rich nucleus pulposus (NP); a peripheral, fibrocartilaginous annulus fibrosus (AF) with a highly ordered, cross-ply lamellar structure; and superiorly and inferiorly two cartilaginous end plates that interface with the adjacent vertebrae [5]. These three structures act together to facilitate the even distribution of compressive loads between the vertebrae, and complex mobility of the intervertebral joint [5]. Disc degeneration is a slowly progressing, cell-mediated cascade that is closely linked to aging, and which ultimately leads to structural and functional derangement of the entire intervertebral joint [2, 4]. Current treatments for disc degeneration and associated low back pain, including both conservative approaches (such as pain medication and physical therapy) and surgery (such as spinal fusion), target symptoms but fail to restore healthy disc structure and function, and often have poor long-term efficacy [4, 6–8].

Degeneration manifests initially in the NP, where an inflammation-mediated reduction in proteoglycan content and hydration compromises resistance to compressive loads [4, 5, 9, 10]. Cell-based strategies to treat disc degeneration by regenerating NP tissue hold significant promise, but key challenges remain [11, 12]. One of these is the inability of therapeutic cells to effectively mimic the performance of native NP cells, which exhibit a unique ability to survive in the oxygen and nutrient-poor disc microenvironment whilst simultaneously secreting large quantities of proteoglycan-rich extracellular matrix (ECM) [5]. Unique amongst skeletal cell types, NP cells arise from the embryonic notochord, a midline structure that serves as both a source of long-range morphogens to guide patterning of the axial skeleton, and as the structure that physically gives rise to the discrete NPs themselves [13–16]. In humans, the rapid and complete disappearance of large, vacuolated, notochordal-like NP cells by skeletal maturity, and their replacement by smaller “mature” NP cells that resemble articular cartilage chondrocytes, is thought to contribute the onset of disc degeneration later in life [5, 17, 18]. In contrast, other species such as mice retain significant numbers of notochordal-like NP cells throughout adulthood and do not typically experience age-associated disc degeneration analogous to humans [18, 19]. Due to this retention of notochordal-like NP cells, mice represent an excellent model with which to study dynamic cell heterogeneity within the postnatal NP. Elucidating the molecular mechanisms underlying the phenotypic transition from notochordal to mature, chondrocyte-like NP cells is important to inform the development of more effective stem cell-based therapeutics, by establishing the optimal characteristics for *de novo* elaboration of the aggrecan-rich NP ECM [20]. Indeed, there is emerging evidence that reprogramming cells to mimic the developmental phenotypes of NP cells may enhance their regenerative potential [21].

Previously, we used bulk RNA-sequencing studies to demonstrate that the function of notochord-derived NP cells transitions from one focused on embryonic patterning to one focused on growth and ECM deposition [22]. We hypothesized that these bulk gene expression changes are due to emergent NP cell subpopulations that drive deposition of the proteoglycan-rich ECM necessary to sustain disc mechanical function in adulthood. To test this hypothesis, the current study used single cell RNA sequencing (scRNA-seq) and identified an emergent NP cell subpopulation in the postnatal mouse disc that exhibits high expression of NP-specific ECM genes and the surface marker Cd9. Additionally, we used a goat model to show that the relative number of these Cd9+ cells decreases significantly coincident with disc degeneration, supporting their association with the maintenance of the healthy NP ECM.

## Results

### Global Single Cell RNA Sequencing Findings

To enrich for notochord-derived NP cells prior to scRNA-seq, we used the shh-cre:R26R-tdTomato (shh/tdTomato) mouse model, which takes advantage of the fact that all cells of the embryonic notochord express the morphogen sonic hedgehog. This generates a fate map whereby notochord-derived cells and their progeny constitutively express tdTomato. NP-specific tdTomato expression was confirmed through *in situ* fluorescence imaging of lumbar discs from birth to postnatal day (P)60, which demonstrated that tdTomato was confined to the NP (**Figure 1A**). Immunofluorescent staining on the same histological sections demonstrated that putative NP cells co-expressed the NP specific marker cytokeratin 19 (Krt19) [23] at each age examined, confirming their identity. For single cell RNA sequencing, cells were isolated through gross dissection of discs from caudal, lumbar and thoracic spines pooled from litter-matched P30 shh/Tomato mice, followed by brief collagenase digestion and fluorescence-activated cell sorting (**Figure 1B**). Three complete replicate scRNA-Seq experiments were performed and the results were pooled prior to analysis.

**Figure 1.**
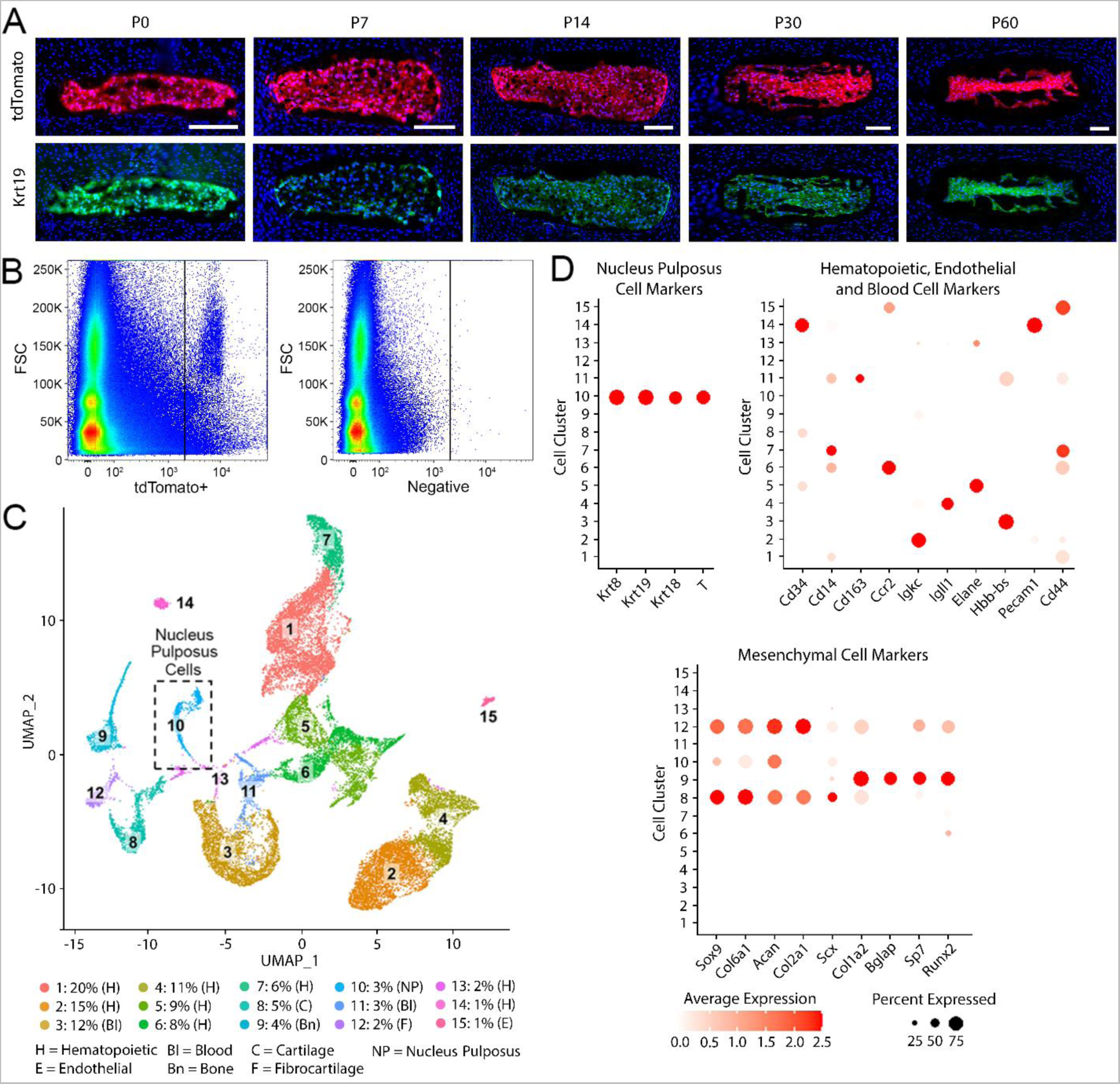
**A.** In situ fluorescence imaging of sections from shh/Tomato mouse lumbar spines showing tdTomato+ cells (red) localized to the NP throughout postnatal growth. Krt19 immunofluorescence (green) on the same sections was used to confirm NP cell identity. Blue: cell nuclei (DAPI); mid-sagittal sections; scale = 100µm. **B.** Representative plots from fluorescence activate cell sorting used to enrich tdTomato+ cells. **C.** UMAP plot of scRNA-seq results of cells isolated from P30 mice showing clustering of 15 different cell populations. **D.** Dot plots showing cluster-specific expression of NP cell markers, hematopoietic, endothelial and blood cell markers, and mesenchymal cell markers.

After quality control, a total of 27066 cells with a median of 2311 genes/cell and a median of 11660 unique molecular identifiers (UMIs)/cell were identified. Unsupervised clustering identified 15 unique cell subpopulations (**Figure 1C**). Surprisingly, despite enriching for tdTomato+ cells, the sequenced cell population was heterogenous and included significant numbers of hematopoietic and mesenchymal lineage cells identified through cluster-specific expression of established markers (**Figure 1D**). This unexpected heterogeneity may be due to leakage and transfer of tdTomato protein from notochordal cells to adjacent cell populations, which is detectable by the laser during cell sorting even when present at low levels. Nucleus pulposus cells, identified through cluster-specific expression of cytokeratins (Krt) 8, 18 and 19, and brachyury (T) comprised 3% of the total cell population (**Figure 1D**).

### Identification of Nucleus Pulposus Cell Subpopulations

Using the above NP marker genes, a total of 936 notochord-derived NP cells with a median of 1445 genes/cell and a median of 4854 UMIs/cell were identified within the total sequenced cell population. Focusing solely on these NP cells, we repeated unsupervised clustering, identifying two distinct NP cell subpopulations (clusters 1 and 2, comprising 76% and 24% of NP cells, respectively, **Figure 2A**). Differentiation trajectory analysis was used to stratify these subpopulations, with cells in clusters 1 and 2 aligning along a pseudo-timeline (**Figure 2B**). Additionally, RNA velocity analysis demonstrated that the direction of differentiation was predominantly towards cluster 2 (**Figure 2C**). Based on this relationship, cells in clusters 1 and 2 were labeled “early stage” and “late stage” NP cells, respectively.

**Figure 2.**
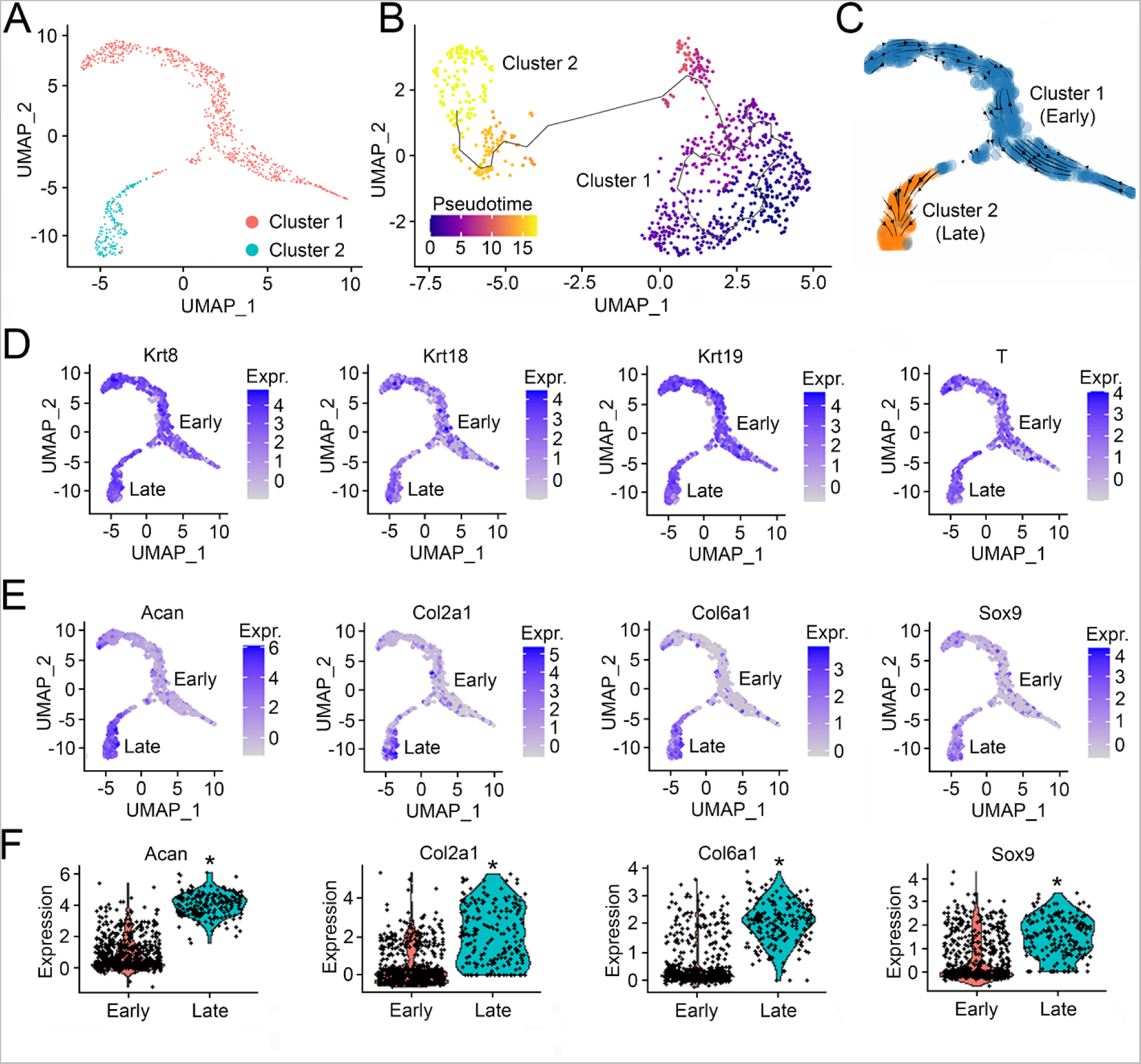
**A.** UMAP plot of scRNA-seq results for NP cells only, identified two distinct cell subpopulations. **B.** Differentiation trajectory analysis demonstrated that NP cells in clusters 1 and 2 aligned along a pseudo timeline; and **C.** RNA velocity analysis showed that the direction of differentiation was predominantly from cluster 1 to cluster 2. Based on these findings, cluster 1 and 2 cells were denoted “early” and “late” stage NP cells, respectively. **D.** UMAP plots for NP marker genes, showing that Krt8, Krt18, Krt19 and T were expressed across both early and late-stage NP cells. **E.** UMAP plots and **F.** Violin plots showing significantly (*) higher expression of NP-specific ECM genes in late-stage NP cells.

There were a total of 317 genes that exhibited significantly different expression levels in late versus early stage NP cells. Of these, 274 genes were upregulated (**Supplementary Table 1**) and 43 genes were downregulated (**Supplementary Table 2**). With respect to NP specific markers Krts 8, 18 and 19, and T, these were expressed at similar levels across both cell clusters (**Figure 2D**), with the exception of Krt18, which was modestly but significantly (0.61 log2 fold) higher in late-stage NP cells. Amongst the top upregulated genes in late stage NP cells were those encoding established components of the healthy NP ECM (**Figures 2E and F**), including aggrecan (Acan, 4.11 log2 fold higher), collagen II-alpha 1 (Col2a1, 3.25 log2 fold higher) and collagen VI-alpha 1 (Col6a1, 2.2 log2 fold higher), in addition to the master chondrogenic transcription factor Sox9 (1.33 log2 fold higher). Cell cycle analysis demonstrated that the majority of both early and late-stage NP cells were in the G1 (resting) phase, and expression of genes related to cell proliferation was modest or absent in both populations (**Supplementary Figure 1**).

### Pathway Analysis

To provide mechanistic insights into the relationship between late and early stage NP cells, we performed gene ontology (GO), Kyoto Encyclopedia of Genes and Genomes (KEGG), and protein-protein interaction (PPI) network analyses. Top GO terms enriched in late versus early-stage NP cells included those related to ECM organization, connective tissue development, and chondrocyte differentiation (**Figure 3A**). With respect to KEGG analysis, we identified 5 pathways that were enriched in late versus early-stage NP cells: ECM-receptor interaction, protein digestion and absorption, PI3K-Akt signaling, focal adhesion, and transforming growth factor-beta (TGF-β) signaling (**Figures 3B, C** and **D**, and **Supplementary Figure 2**). Protein-protein interaction (PPI) network analysis demonstrated significant interaction between elements of these pathways (**Figure 3E**). Genes with the greatest (>5) number of interactions included fibronectin-1 (Fn1), Col6a1, Col6a2, Acan, Col2a1, collagen XI-alpha 1 (Col11a1), Fosb and early growth response factor 1 (Egr1). Using the CellChat tool [24], a total of 93 interactions between and within early and late stage NP cells were identified (**Figure 3F and Supplementary Table 3**). Interactions categories included secreted signaling, ECM-receptor and cell-cell contact. With respect to secreted signaling, there were 7 ligand-receptor interactions from early stage to late stage NP cells, 2 from late stage to early stage NP cells, and 22 from late stage to late stage NP cells (**Figure 3F**). The secreted signaling interactions from early to late stage NP cells involved either pleiotrophin (Ptn) or osteopontin (Spp1).

**Figure 3.**
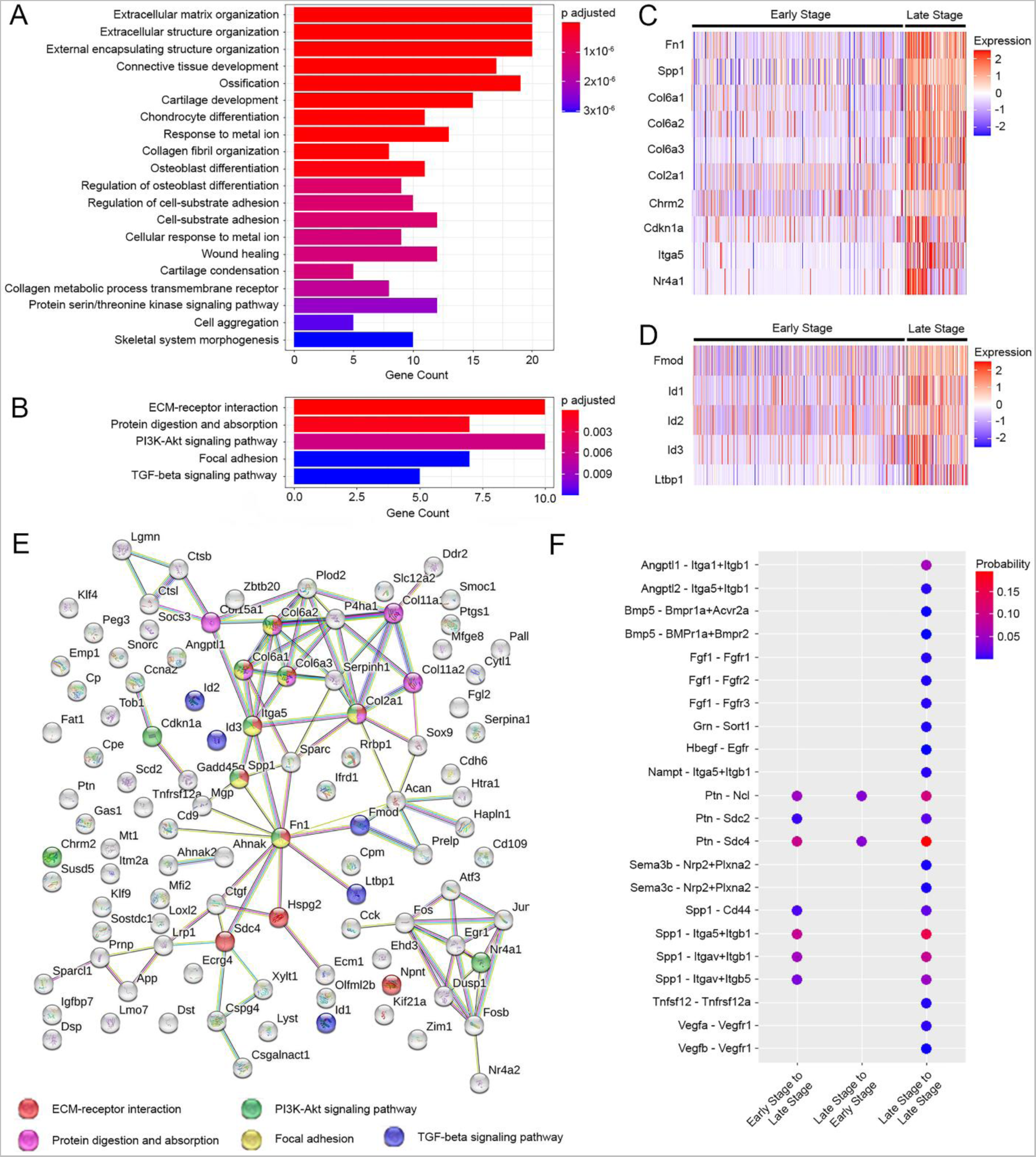
Pathway analysis. **A.** Top 20 GO terms significantly enriched in late versus early-stage NP cells. **B.** Pathways significantly enriched in late versus early-stage NP cells identified by KEGG analysis. Heatmaps showing differentially expressed genes in the **C.** PI3kT-Akt and **D.** TGF-β signaling pathways, respectively, between late and early stage NP cells. **E.** Gene network analysis for all 5 pathways enriched in late versus early stage NP cells. **F.** Secreted signaling interactions between NP cell populations identified using CellChat.

### Surface Marker Expression and Spatial Localization of NP Cell Subpopulations

To facilitate *in situ* spatial localization of distinct NP cell subpopulations within postnatal mouse discs, we identified two candidate surface markers in our single cell sequencing results, Cd109 and Cd9, that exhibited significantly higher expression levels in late versus early-stage NP cells (2.56 and 1.90-log2 fold higher, respectively; **Figures 4A** and **B**, and **Supplementary Table 1**). Immunofluorescent histology demonstrated that Cd109 protein expression was relatively homogeneous throughout the entire NP at all ages examined, with staining at the periphery being marginally more intense (**Figure 4C**). In contrast, spatially heterogenous protein expression was striking for Cd9, with immunopositive cells being localized to the periphery of the NP (**Figure 4C**). Subjectively, the relative number of these cells became progressively greater with increasing age from P0 to P90. This observation was confirmed using flow cytometry and double immunolabeling for Krt19 and Cd9, which demonstrated that the number of Cd9+ NP cells as a percentage of total NP cells did indeed increase with increasing age, from less than 1% at P0 to more than 30% at P90 (**Figure 4D**). Next, we examined ECM elaboration during postnatal growth using Alcian blue (for GAG) and picrosirius red (collagen) staining (**Figure 4E**). Glycosaminoglycan deposition was spatially heterogenous, appearing as a halo surrounding a central cluster of cells that increased in size with increasing age and mirroring the emergence of Cd9+ NP cells (**Figure 4E**).

**Figure 4.**
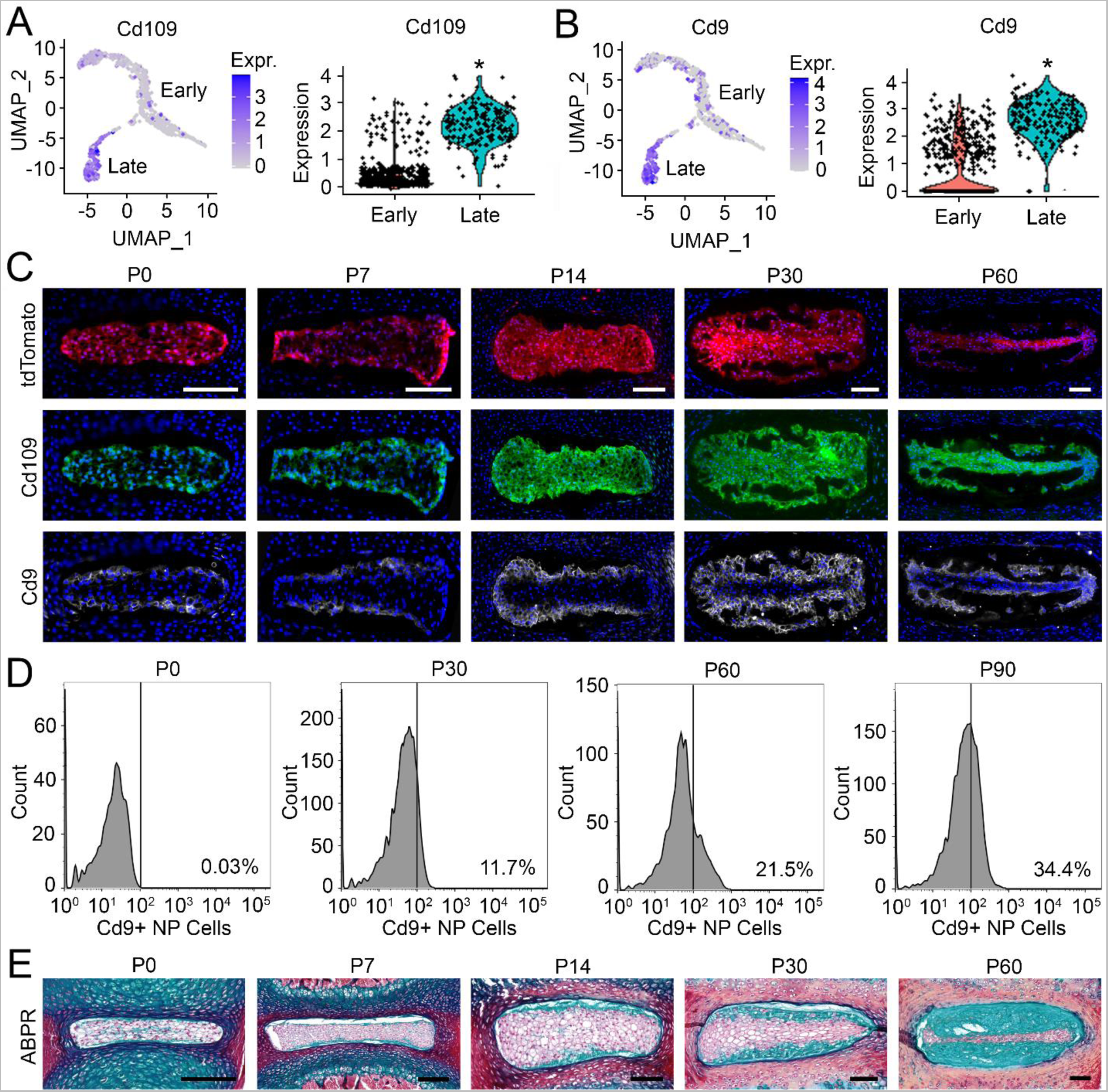
UMAP and violin plots showing significantly (*) higher expression of the surface markers **A.** Cd109 and **B.** Cd9 in late versus early stage NP cells. **C.** *In situ* fluorescence imaging of tdTomato+ cells (red), and corresponding immunofluorescence imaging of Cd109+ (green) and Cd9+ (white) cells in the NPs of mouse lumbar spines during postnatal growth. Cd9+ cells were localized to the NP periphery, with relative numbers increasing with postnatal age. Blue (DAPI) = cell nuclei; Midsagittal sections; 100µm. **D.** Flow cytometry analysis showing relative increases in the number of tdTomato/Krt19/Cd9+ positive cells with increasing postnatal age. **E.** Representative sections showing progressive accumulation of glycosaminoglycan-rich ECMs at the NP periphery with increasing postnatal age. Alcian blue and picrosirius red staining; mid-sagittal sections; scale = 100µm.

To further investigate whether Cd9 was a marker of the healthy NP cell phenotype, we used a goat model of moderate severity disc degeneration. In this model, 1U of the enzyme chondroitinase ABC was injected into the NP of goat lumbar discs, resulting in moderate severity degeneration after 12 weeks (**Figure 5A**). Successful induction of degeneration was confirmed postmortem via semi-quantitative histological grading (**Figure 5B**). Using immunohistochemistry, we confirmed the presence of Cd9+ NP cells in both healthy and degenerate goat discs (**Figure 5C**); however, the number of Cd9+ cells as a percentage of total NP cells was significantly lower in degenerate discs (**Figure 5D**).

**Figure 5.**
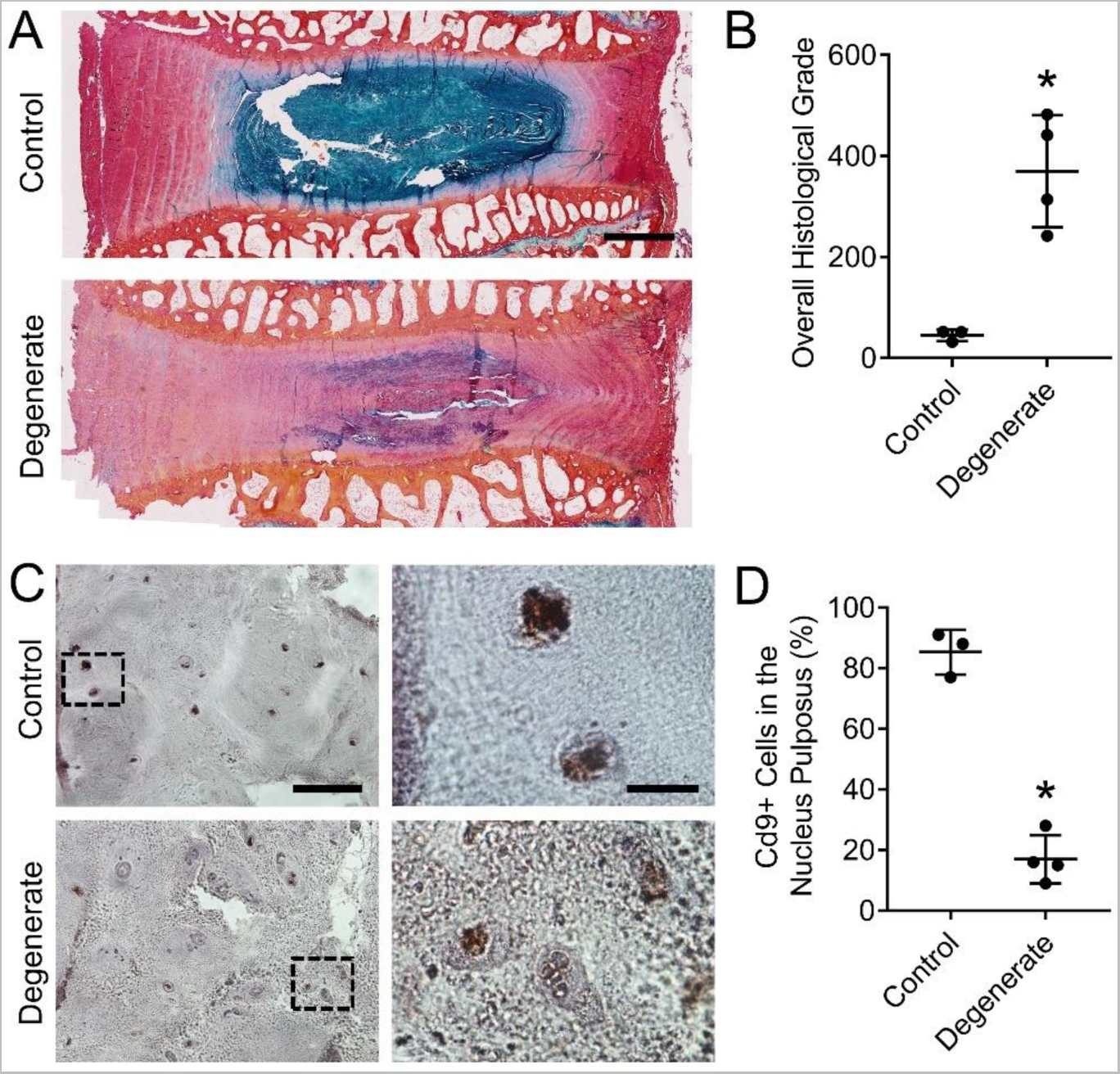
**A.** Representative histological sections of healthy and degenerate goat lumbar discs. Alcian blue and picrosirius red-staining; mid-sagittal sections; scale = 2mm. **B.** Semi-quantitative grade of healthy and degenerate goat discs. *p<0.05 vs healthy; N=3-4; Students t-test. **C.** Representative immunostaining of Cd9+ cells in the NPs of healthy and degenerate goat discs. Mid-sagittal sections; scale = 100µm (left) and 20µm (right). **D.** Quantification of the relative numbers of Cd9+ cells in the NPs of healthy and degenerate goat discs. *p<0.05 vs healthy; N=3-4; Students t-test.

## Discussion

The notochordal origin of NP cells is unique amongst skeletal cell types. The notochord is a rod-like structure that is present embryonically in all chordates, and functions as a source of long-range morphogens that regulate patterning of the axial skeleton. In mice, beginning at around embryonic day E12.5, the notochord undergoes a transformation, contracting within the regions of the future vertebrae, and expanding within the regions of the future discs, to ultimately form the NPs. This process occurs similarly across other vertebrates, including humans. The notochordal origin of all NP cells in the mouse disc throughout postnatal growth and adulthood has been confirmed though fate-mapping studies that have leveraged notochordal markers such as Shh [13], Noto [15], and FoxA2 [25]. A recent study also showed that all NP cells express the cytoskeletal marker Krt19 throughout life [26]. We confirmed this in the current study, and leveraged expression of Krt19 and other NP-specific markers such as Krt8 and 18, and brachyury (T) to identify NP cells within our overall sequenced cell population.

The eventual depletion of early stage (historically referred to as “notochordal” cells) in other species, including humans, occurs for reasons still not well understood, and may be due to both longer life span and species-specific physical and biochemical microenvironmental factors. This loss of notochordal NP cells in humans is considered to be a predisposing factor in the initiation and progression of disc degeneration later in life, as notochordal NP cells and their secretome have been shown to be pro-regenerative properties, enhancing proliferation and ECM production, and reducing inflammation [27, 28]. In this study we used scRNA-seq to demonstrate emergent heterogeneity amongst notochord-derived NP cells in the postnatal mouse disc (shown schematically in **Figure 6**).

**Figure 6.**
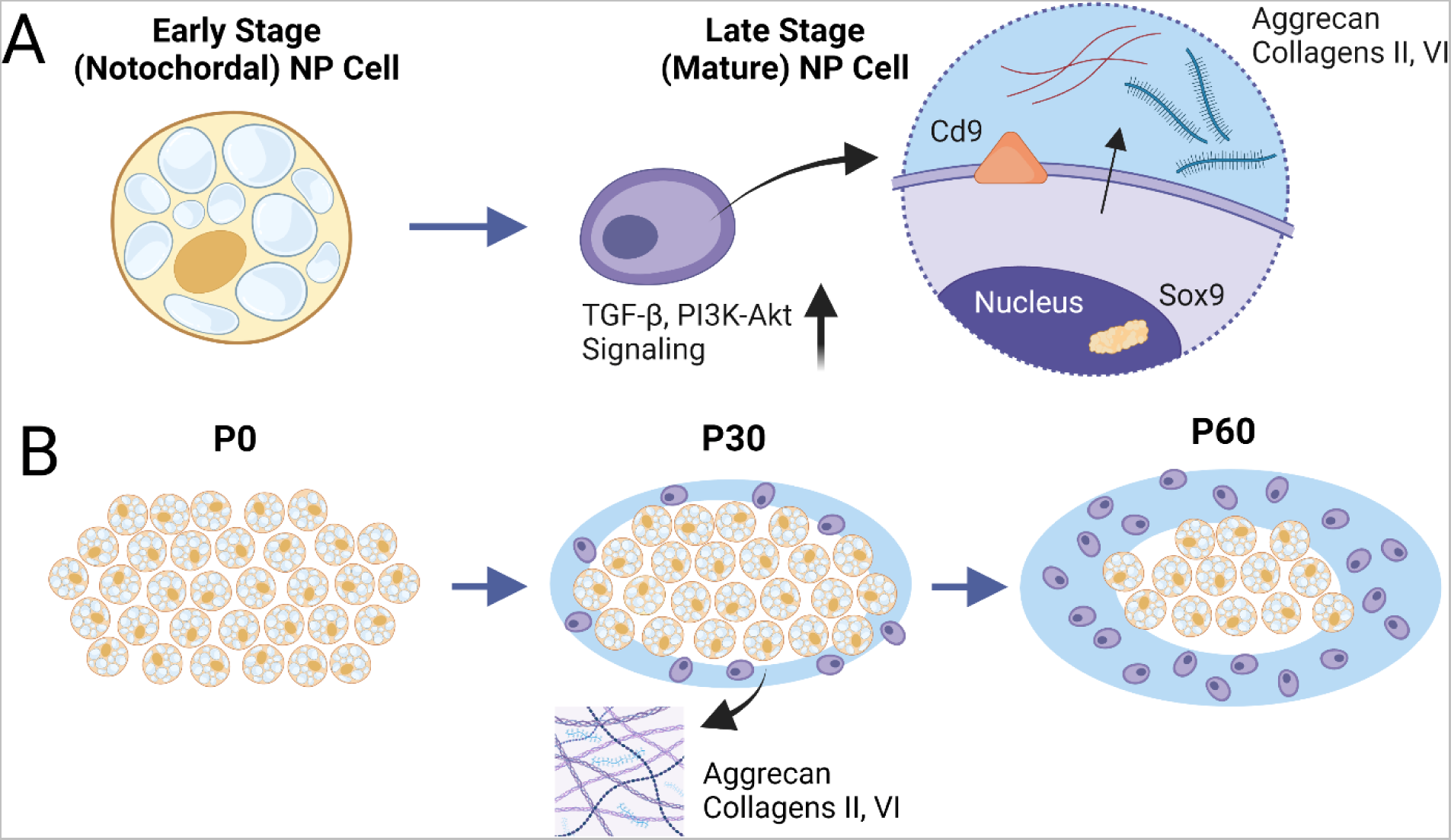
Schematic representations of **A.** the transition from early to late-stage NP cells, which is characterized by increased TGF-β and PI3k-Akt signaling, and expression of aggrecan, collagens II and VI, and Cd9; and **B.** emergent late-stage NP cells during postnatal growth with concomitant peripheral ECM deposition. Created with BioRender.com, individual license AG25DQC3X0.

Specifically, we established the existence of early and late-stage NP cells, which likely correspond to “notochordal” and “mature” NP cells, respectively, consistent with our study hypothesis. These cells exhibited distinct gene expression profiles reflecting unique functional roles. Specifically, our findings suggest that early stage, notochordal NP cells residing at the center of the NP serve as progenitors that give rise to late stage, mature NP cells that reside at the periphery. Histological findings suggest that this transition from early to late-stage NP cells occurs progressively during postnatal growth, and may ultimately lead to the early stage progenitor population being exhausted. A recent study described the emergence of notochord-derived, chondrocyte-like cells in the postnatal mouse disc from 16 months-of-age [26]. These cells, also termed late stage NP cells, were observed to fuse into nest-like structures and were associated with pathological aging. In contrast, the late stage cells identified in the current study were emerged early during postnatal growth, and expressed anabolic markers typically associated with the healthy NP ECM, including aggrecan, collagen II and sox9. Our results suggest that these cells are primarily responsible for elaborating the proteoglycan-rich ECM of the healthy NP, and may represent an intermediate phenotype that transitions towards an aged or pathological state later in life.

In the current study, pathway analysis of scRNA-seq results identified both TGF-β and PI3K/Akt signaling as significantly enriched in late versus early stage NP cells. Embryonically, TGF-β signaling has been shown to be important for formation of both the NP and annulus fibrosus [29, 30]. In a prior study we showed that TGF-β expression is elevated in mouse NP cells at the onset of postnatal growth concomitantly with elevated ECM expression [22]. *In vitro*, adult NP cells treated with exogenous TGF-β increase production of healthy ECM components [31]. In the current study, elevated TGF-β signaling in late stage NP cells, specifically, is consistent with a central role in ECM elaboration and organization. PI3K/Akt signaling is a complex pathway involving more than 150 proteins, is activated by several growth factors, cytokines, and mechanical stimuli, and performs functions in many cellular processes essential for homeostasis, including regulation of cell proliferation, survival, inflammation, metabolism, and apoptosis [32, 33]. To our knowledge, PI3K/Akt signaling has not been studied in the context of disc development, though inhibition of PI3K/Akt signaling in rat NP cells was found to decrease both proliferation and ECM expression [34].

Evidence from prior studies suggests that secreted factors from early stage (notochordal) NP cells regulates ECM production in late stage (mature) NP cells [28]. In the current study, analysis of cell-cell interactions identified two secreted factors, Ptn and Spp1, that may signal from early to late stage NP cells. Pleiotrophin is a developmentally-regulated heparin-binding protein abundant in fetal, but not adult articular cartilage, which has been shown to inhibit proliferation and stimulate glycosaminoglycan synthesis in mature chondrocytes [35]. Interestingly, a prior study found that Ptn is expressed at higher levels in NP cells compared to articular chondrocytes [36]. It is possible that secretion of Ptn by early stage NP cells stimulates proteoglycan production by late stage NP cells. Osteopontin is a secreted, integrin-binding, glycosylated phosphoprotein important in biomineralization and pathological cartilage remodeling [37]. Additional roles include regulating cell migration and adhesion, and ECM organization [38], with Spp1 knockout mice exhibiting aberrant wound healing [39]. Interestingly, Spp1 was previously shown to be expressed in the notochord [40, 41], and a very recent study used single cell sequencing analysis to identify Spp1 as playing a key role in intercellular crosstalk during early human disc formation [41].

We also identified two surface markers, Cd9 and Cd109, that were significantly elevated in their expression in late versus early stage NP cells. Two recent studies used proteomics to identify Cd109 as a novel NP-specific marker in both mice and humans [42, 43]; however, to our knowledge, no previous studies have examined the role of Cd9 in the NP. Interestingly, we found that Cd109 expression was relatively homogenous across all tdTomato+ NP cells, while Cd9 expression was clearly confined to a subset of NP cells that were localized to the peripheral zone. Further, the relative number of Cd9+ cells increased with age, and was co-incident with the emerging halo of glycosaminoglycan-rich ECM. These findings suggest that Cd9 may be a more faithful marker of late-stage NP cells, specifically, relative to Cd109. As further evidence that these cells are associated with the healthy NP ECM, we used a preclinical goat model to show that the number of Cd9+ NP cells significantly diminished with disc degeneration. Cd9, a member of the tetraspanin family of molecules, is a cell surface glycoprotein expressed by many different cell types, and which has diverse roles including in growth, differentiation, cell adhesion, motility and regulation of inflammation [12]. Interestingly, in other cell types including immune, lung and endothelial cells, Cd9 interacts with integrins, reflecting a key role in cell adhesion and interactions with the ECM [44]. In keratinocytes, Cd9, through interactions with E-cadherin, has been shown to activate PI3K-Akt signaling (enriched in late-stage NP cells) [45]. Notably, Cd9 is also marker of extracellular vesicles [46].

This study had a number of limitations. Despite using shh/tdTomato mice to enrich for notochord-derived cells, our sequenced cell population was heterogenous, and included both mesenchymal and hematopoietic cells. This heterogeneity was surprising given *in situ* imaging that showed tdTomato expression confined to the NP region. Similar heterogeneity was reported previously for cells enriched from other tdTomato reporter mice [47]. As notochordal cells are highly secretory, it is possible that tdTomato protein is transferred to adjacent cell populations and is detectable by the laser during cell sorting even when present at low levels. This contention is supported by previous in vitro co-culture studies of progenitor and differentiated cells, where chemical fluorescent markers are readily transferred from labeled to adjacent unlabeled cells [48]. While the NP cell cluster was clearly distinguishable within the broader cell population through specific expression of established NP marker genes, this heterogeneity meant that the overall number of NP cells available for analysis was relatively small. Experimental outcomes in this study were limited to gene expression and *in situ* validation assays, and while our findings highlight potential functional differences between early and late-stage NP cells, these must now be confirmed through *in vivo* genetic manipulation of these cell subpopulations during disc development and *in vitro* cell culture experiments.

In conclusion, in this study, we provide novel insights into the fate and function of different cell types that comprise the postnatal NP. Specifically, we showed that early stage NP cells give rise to late stage NP cells that are characterized by higher expression of healthy ECM genes and the cell surface marker Cd9. Improved understanding of the developmental mechanisms underlying regulation of ECM deposition in the postnatal NP may inform improved regenerative strategies for disc degeneration and associated low back pain.

## Experimental Procedures

### Animals and Study Design

All animal studies were carried out with approval from the Institutional Animal Care and Use Committees of the University of Pennsylvania and/or the Corporal Michael J Crescenz Philadelphia VA Medical Center. To obtain notochord-derived NP cells for scRNA-Seq and for *in situ* validations experiments, we used a Shh/TdTomato mouse model, which leverages the fact that all cells of the embryonic notochord express Shh, and produces constitutive expression of tdTomato at the ROSA26 locus in these cells and their progeny, including when Shh is no longer expressed. Shh/TdTomato mice were generated by breeding Shh:cre mice (JAX005562; Jackson Laboratory, Bar Harbor, ME, USA) [22] with R26R:tdTomato mice (JAX007905; Jackson Laboratory). Animals were euthanized via carbon dioxide inhalation and spines were harvested for the experiments described.

To examine how the number of Cd9+ NP cells changes as a function of disc degenerative condition, we used an established goat model of lumbar disc degeneration. Moderate severity degeneration was induced in the lumbar discs (L1-2 to L4-5) of six, 2-year-old large frame goats via injection of 1U chondroitinase ABC (ChABC) into the NP as described previously [49]. The detailed surgical procedure was previously published. Briefly, under general anesthesia, the lumbar spine was exposed using an open, lateral, retroperitoneal transpsoatic approach. ChABC was suspended in 200μl of sterile saline and injected into the NP using a 22G spinal needle under fluoroscopic guidance. Adjacent, non-degenerate disc levels (T13-L1 and L5-6) were maintained as healthy, non-degenerate controls. The incision was closed in layers, and the animal recovered and returned to housing. For analgesia, animals were administered per-operative transdermal fentanyl (2.5mcg/kg/hr) and intravenous flunixin meglumine (Banamine, 1.1mg/kg) for analgesia, while florfenicol (40 mg/kg) was administered for antimicrobial prophylaxis. Animals were housed in groups with unlimited access to exercise, and evaluated daily by a veterinarian for signs of pain, behavior changes or gait abnormalities for the duration of the study. A total of 4 degenerate and 3 healthy control discs were used for this study, with remaining discs allocated to other, unrelated studies. Twelve weeks after inducing degeneration, animals were euthanized via an overdose of sodium pentobarbital and lumbar spines harvested for the experiments described below.

### Cell Isolation, Single Cell RNA-Sequencing and Analysis

Spinal columns were excised from mice postmortem. Under a dissecting microscope, dorsal boney elements and the spinal cord were removed leaving the isolated ventral spinal column comprising the vertebrae and discs. Whole discs in the lumbar, thoracic and caudal regions were carefully isolated by cutting through the vertebral boney endplates with a scalpel, and bisected to expose the NPs. Isolated discs were placed in a phosphate buffered saline containing 25 mM HEPES and 0.1% FBS, finely diced, then digested in collagenase (1ml/mg; Sigma-Aldrich, St Louis, USA) for 30 mins. After digestion, cells were strained through a 50 µm filter. TdTomato+ cells were then enriched using fluorescence activated cell sorting [22].

Single cell sequencing of enriched TdTomato+ cells was performed using methods similar to those described previously [50]. The experiment was performed in triplicate. Viability (~95%) was confirmed using trypan blue staining, and three batches of single cell libraries were generated. Cells were loaded in the Chromium Controller (V3 chemistry version; 10X Genomics, San Francisco, USA), barcoded and purified according to the manufacturer’s instructions, and 2×150 bp paired-end sequencing was performed at a depth of approximately 400 million reads (HiSeq; Illumina, San Diego, USA). Cellranger (Version 3.0.2, 10 Genomics) was used to demultiplex reads, followed by extraction of cell barcode and unique molecular identifiers (UMIs), and the cDNA insert was aligned to a modified reference mouse genome (mm10). Poor quality cells (< 200 genes or >5% mitochondrial genes) and doublets (> 6000 genes) were excluded. Gene expression was natural long transformed and normalized using the Seurat v4 toolkit and R Statistical Software (v4.1.2; R Core Team 2021) [51].

Results from the three sequencing batches were integrated for analysis. The principal component dimension number was set as 50, with 2000 highly variable genes. To identify major cell types, the FindAllMarkers code statement was used to label single cells. Unsupervised clustering was initially performed for the entire sequenced cell population and visualized using Uniform Manifold Approximation and Projection (UMAP) plots. Cell populations were identified using known markers for NP, mesenchymal, and hematopoietic, endothelial and blood cell markers. Unsupervised clustering was subsequently repeated for the cell subpopulation identified as NP cells and visualized using UMAP plots. Significant differences in gene expression between identified subpopulations of NP cells were determined. Cell cycle analysis was performed for NP cell subpopulations using previously defined cell cycle phase marker genes [52] in the CellCycleScoring package. Cells in the G1 phase were classified as resting, while those in the G2M or S phases were classified as proliferating. Differentiation trajectory analysis was performed for NP cell subpopulations using the Monocle 3 package [53]. The RNA velocity of NP cells in each cluster was calculated using the packages ‘velocity’ and ‘scVelo’ [54]. To determine biological pathways enriched in specific NP cell subpopulations, GO and KEGG analyses were performed using the clusterProfiler (v4.5.3) package for significantly differentially expression genes with a log2 fold change ≥1. Protein-Protein Interaction Network Construction and Module Analysis was used to obtain interaction relationships among differentially expressed genes [55], with the minimum required interaction score set to high confidence (0.7). Examination of cell-cell interactions within and between different NP cell subpopulations was performed using the CellChat tool [24]. Using the paracrine/autocrine signaling interaction dataset of CellChatDB as the referencing database, the communication probability was computed using a truncated mean of 20%. Cell-cell communication was subsequently inferred and the cell-cell communication network was aggregated with default parameters. The number of interactions was visualized to show the aggregated cell-cell communication network and signaling from each cell cluster.

### Histology

Lumbar spines (vertebral columns with posterior boney elements removed) were isolated postmortem from shh/tdTomato mice aged 0, 7, 14, 30 and 60 days (n=3 per age), fixed in buffered 10% formalin overnight and decalcified in formic and ethylenediaminetetraacetic acids for 24 hours (Formical 2000; StatLab Medical Products, McKinney, USA) for 24 hours. For fluorescent and immunofluorescent localization of NP cells, spines were then immersed into 20% sucrose and 2% polyvinylpyrrolidone (PVP) at 4°C overnight, embedded in OCT compound (Sakura), and mid-sagittal 10 μm sections were cut on a freezing microtome (NX-70 Cryostar, Thermo Fisher Scientific Inc, Waltham, MA). For immunofluorescence, sections were incubated with primary antibodies against rabbit anti-Krt19 (1:200; NB100687; Novus Biologicals, Centennial, USA), rat anti-Cd9 (1:200; ab82390; Abcam Inc) or rabbit anti-Cd109 (1:200; ab203588, Abcam Inc) at 4°C overnight, followed by Alexa Fluor 488-conjugated donkey anti-rabbit (1:1000; A21206; Invitrogen,) or Alexa Fluor 647 goat anti-rat (1:1000, Invitrogen, A21247) secondary antibodies for 1 hour at room temperature. For negative controls, primary antibodies were omitted. Finally, sections were counterstained with DAPI (Fluoromount-G; SouthernBiotech; Birmingham, USA) and imaged using a confocal microscope (LSM 9080; Carl Zeiss AG; Oberkochen, Germany). For histochemical assessment of progressive ECM deposition in mouse discs, lumbar spines (n=3 per age) were fixed and decalcified as above, and then processed into paraffin. Mid-sagittal, 10 μm sections were cut and double stained with Alcian blue and picrosirius red for qualitative assessment of glycosaminoglycan and collagen distribution, respectively, and imaged using bright field microscopy (Eclipse 90i; Nikon, Tokyo, Japan).

For goat samples, vertebra-disc-vertebra segments were fixed in 10% formalin, completely decalcified in formic and ethylenediaminetetraacetic acids, and processed into paraffin. For semi-quantitative grading of histological condition, mid-sagittal, 8μm-thick sections were cut and double stained with Alcian blue (glycosaminoglycans) and picrosirius red (collagen). Semi-quantitative histological grading was performed by 3 blinded assessors (averaged) as described previously [49]. Parameters assessed included NP cellularity, NP ECM, endplate structure, NP-AF boundary, and AF organization, each on a continuous scale from 0 (best) to 100 (worst). An overall grade (ranging from 0 to 500) was calculated as the sum of these individual scores. For immunohistochemical assessment of Cd9 expression on goat sections, 8 µm sections were rehydrated, and antigen retrieval performed at 60°C in a Tris-EDTA pH 8.0 buffer, overnight. Endogenous peroxidase activity was blocked by treating sections with 3% hydrogen peroxide for 12 minutes, followed by Background Buster (Innovex Biosciences; Richmond, USA) for 10 minutes at room temperature to block non-specific protein binding. Sections were then incubated with rabbit anti-Cd9 (Abcam) at 4°C overnight. Staining was visualized using the Vectastain Elite ABC-Peroxidase Kit (Vector Laboratories; Burlingame, USA) and diaminobenzidine chromogen (Thermofisher Scientific; Waltham, USA) according to the manufacturer’s protocol. Finally, sections were counterstained with hematoxylin QS (Vector Laboratories; Burlingame, USA) and cover-slipped with aqueous mounting medium (Agilent; Santa Clara, USA). For analysis, all slides were imaged under bright field light microscopy. Three randomly selected regions of interest within the NP were analyzed. For each region, the number of immunopositive cells was counted and normalized as a percentage of the total number of cells present. Cell counting for each region was performed by three individuals who were blinded to the study groups and averaged across scorers prior to statistics.

### Flow Cytometry

Notochord-derived NP cells were enriched from shh/tdTomato mice (cells from 6 animals pooled per timepoint) aged 0, 30, 60 and 90 days as described. Cells were stained sequentially with a rabbit polyclonal Krt19 antibody conjugated to Alexa Fluor 647 (1:200; catalog number NB100-687AF647; Novus Biologicals) and a rat monoclonal Cd9 antibody conjugated to Alexa Fluor 750 (1:200; catalog number NBP1-44876AF750; Novus) for 45 minutes on ice. Cells were then washed twice in phosphate buffered saline containing 2% fetal bovine serum (ThermoFisher Scientific; Waltham, MA) and analyzed using a flow cytometer (LSRII; BD Biosciences; Franklin Lakes, USA). NP cells were identified within the total population as double positive for tdTomato and Krt19, and Cd9+ cells as a percentage of this NP cell population at each age were determined using Flowjo software (BD Biosciences). Unstained cells served as negative controls.

### Statistical Analyses

Statistically-significant differences in histological grade of degeneration and the percentage of NP cells immunopositive for Cd9 between healthy and degenerative goat discs were established using Student’s t-tests using Prism software (GraphPad Software, San Diego, CA). Data is presented as mean ± standard deviation and significance was defined as p<0.05.

## Acknowledgements

Funding for this work was received from the National Institutes of Health (R01AR077435, R21AR077261 and P30AR050950) and the Department of Veteran’s Affairs (I01RX001321). Technical support received from staff at the Center for Applied Genomics at the Children’s Hospital of Philadelphia is gratefully acknowledged.

## Data Availability Statement

The scRNA-Seq datasets generated and analyzed during this study will be made available in the NCBI Gene Expression Omnibus (GEO) repository upon manuscript acceptance.

Additional study data will be made available upon reasonable request to the corresponding author.

## Author Contribution Statement

CZ contributed to conceptual design, performed experiments and interpretation of results, and drafted the manuscript; YKL, LZ, MW and LY performed experiments; TPS performed goat surgeries and supervised animal care; RLM contributed to conceptual design and interpretation of results; NRM performed goat surgeries and contributed to conceptual design; LQ contributed to conceptual design and interpretation of results; LJS conceived the study, contributed to interpretation of results, and drafted the manuscript. All authors reviewed and approved the final version of the manuscript prior to submission.

## Supplementary Material

**Supplementary Figure 1**. **A.** UMAP of NP cells showing cell cycle state. **B.** Violin plots showing relative expression of cell proliferation markers in early and late stage NP cells.

**Supplementary Figure 2.** Heatmaps showing differentially expressed genes in the **A.** Protein digestion and absorption; **B.** Focal adhesion; and **C.** ECM-receptor interactions pathways, respectively, between late and early-stage NP cells.

**Supplementary Table 1.** Genes significantly upregulated in late versus early-stage NP cells, ranked in descending order by relative log2 fold change.

**Supplementary Table 2.** Genes significantly downregulated in late versus early-stage NP cells, ranked in descending order by relative log2 fold change.

**Supplementary Table 3**. Cell-cell interactions identified using CellChat.

## References

[1] J.L. Dieleman, R. Baral, M. Birger, A.L. Bui, A. Bulchis, A. Chapin, H. Hamavid, C. Horst, E.K. Johnson, J. Joseph, R. Lavado, L. Lomsadze, A. Reynolds, E. Squires, M. Campbell, B. DeCenso, D. Dicker, A.D. Flaxman, R. Gabert, T. Highfill, M. Naghavi, N. Nightingale, T. Templin, M.I. Tobias, T. Vos, C.J. Murray, US Spending on Personal Health Care and Public Health, 1996-2013, JAMA 316(24) (2016) 2627–2646.

[2] A.J. Freemont, The cellular pathobiology of the degenerate intervertebral disc and discogenic back pain, Rheumatology (Oxford) 48(1) (2009) 5–10.

[3] C. Global Burden of Disease Study, Global, regional, and national incidence, prevalence, and years lived with disability for 301 acute and chronic diseases and injuries in 188 countries, 1990-2013: a systematic analysis for the Global Burden of Disease Study 2013, Lancet 386(9995) (2015) 743–800.

[4] P.P. Raj, Intervertebral disc: anatomy-physiology-pathophysiology-treatment, Pain Pract 8(1) (2008) 18–44.

[5] L.J. Smith, N.L. Nerurkar, K.S. Choi, B.D. Harfe, D.M. Elliott, Degeneration and regeneration of the intervertebral disc: lessons from development, Dis Model Mech 4(1) (2011) 31–41.

[6] J.C. Eck, A. Sharan, Z. Ghogawala, D.K. Resnick, W.C. Watters, 3rd, P.V. Mummaneni, A.T. Dailey, T.F. Choudhri, M.W. Groff, J.C. Wang, S.S. Dhall, M.G. Kaiser, Guideline update for the performance of fusion procedures for degenerative disease of the lumbar spine. Part 7: lumbar fusion for intractable low-back pain without stenosis or spondylolisthesis, J Neurosurg Spine 21(1) (2014) 42–7.

[7] L.G. Hart, R.A. Deyo, D.C. Cherkin, Physician office visits for low back pain. Frequency, clinical evaluation, and treatment patterns from a U.S. national survey, Spine (Phila Pa 1976) 20(1) (1995) 11–9.

[8] C. Zhang, S.H. Berven, M. Fortin, M.H. Weber, Adjacent Segment Degeneration Versus Disease After Lumbar Spine Fusion for Degenerative Pathology: A Systematic Review With Meta-Analysis of the Literature, Clin Spine Surg 29(1) (2016) 21–9.

[9] C.L. Le Maitre, A. Pockert, D.J. Buttle, A.J. Freemont, J.A. Hoyland, Matrix synthesis and degradation in human intervertebral disc degeneration, Biochem Soc Trans 35(Pt 4) (2007) 652–5.

[10] J.P. Urban, S. Roberts, Degeneration of the intervertebral disc, Arthritis Res Ther 5(3) (2003) 120–30.

[11] L.J. Smith, L. Silverman, D. Sakai, C.L. Le Maitre, R.L. Mauck, N.R. Malhotra, J.C. Lotz, C.T. Buckley, Advancing cell therapies for intervertebral disc regeneration from the lab to the clinic: Recommendations of the ORS spine section, JOR Spine 1(4) (2018) e1036.

[12] M. Loibl, K. Wuertz-Kozak, G. Vadala, S. Lang, J. Fairbank, J.P. Urban, Controversies in regenerative medicine: Should intervertebral disc degeneration be treated with mesenchymal stem cells?, JOR Spine 2(1) (2019) e1043.

[13] K.S. Choi, M.J. Cohn, B.D. Harfe, Identification of nucleus pulposus precursor cells and notochordal remnants in the mouse: implications for disk degeneration and chordoma formation, Dev Dyn 237(12) (2008) 3953–8.

[14] L. Lawson, B.D. Harfe, Notochord to Nucleus Pulposus Transition, Curr Osteoporos Rep 13(5) (2015) 336–41.

[15] M.R. McCann, O.J. Tamplin, J. Rossant, C.A. Seguin, Tracing notochord-derived cells using a Noto-cre mouse: implications for intervertebral disc development, Dis Model Mech 5(1) (2012) 73–82.

[16] D.L. Stemple, Structure and function of the notochord: an essential organ for chordate development, Development 132(11) (2005) 2503–12.

[17] R. Cappello, J.L. Bird, D. Pfeiffer, M.T. Bayliss, J. Dudhia, Notochordal cell produce and assemble extracellular matrix in a distinct manner, which may be responsible for the maintenance of healthy nucleus pulposus, Spine (Phila Pa 1976) 31(8) (2006) 873–82; discussion 883.

[18] C.J. Hunter, J.R. Matyas, N.A. Duncan, Cytomorphology of notochordal and chondrocytic cells from the nucleus pulposus: a species comparison, J Anat 205(5) (2004) 357–62.

[19] S.N. Tang, B.A. Walter, M.K. Heimann, C.C. Gantt, S.N. Khan, O.N. Kokiko-Cochran, C.C. Askwith, D. Purmessur, In vivo Mouse Intervertebral Disc Degeneration Models and Their Utility as Translational Models of Clinical Discogenic Back Pain: A Comparative Review, Front Pain Res (Lausanne) 3 (2022) 894651.

[20] H. Choi, Z.I. Johnson, M.V. Risbud, Understanding nucleus pulposus cell phenotype: a prerequisite for stem cell based therapies to treat intervertebral disc degeneration, Curr Stem Cell Res Ther 10(4) (2015) 307–16.

[21] S. Tang, J. Richards, S. Khan, J. Hoyland, D. Gallego-Perez, N. Higuita-Castro, B. Walter, D. Purmessur, Nonviral Transfection With Brachyury Reprograms Human Intervertebral Disc Cells to a Pro-Anabolic Anti-Catabolic/Inflammatory Phenotype: A Proof of Concept Study, Journal of orthopaedic research : official publication of the Orthopaedic Research Society 37(11) (2019) 2389–2400.

[22] S.H. Peck, K.K. McKee, J.W. Tobias, N.R. Malhotra, B.D. Harfe, L.J. Smith, Whole Transcriptome Analysis of Notochord-Derived Cells during Embryonic Formation of the Nucleus Pulposus, Scientific reports 7(1) (2017) 10504.

[23] S. Mohanty, R. Pinelli, C.L. Dahia, Characterization of Krt19 (CreERT) allele for targeting the nucleus pulposus cells in the postnatal mouse intervertebral disc, J Cell Physiol 235(1) (2020) 128–140.

[24] S. Jin, C.F. Guerrero-Juarez, L. Zhang, I. Chang, R. Ramos, C.H. Kuan, P. Myung, M.V. Plikus, Q. Nie, Inference and analysis of cell-cell communication using CellChat, Nat Commun 12(1) (2021) 1088.

[25] T.Y.K. Au, T.K. Lam, Y. Peng, S.L. Wynn, K.M.C. Cheung, K.S.E. Cheah, V.Y.L. Leung, Transformation of resident notochord-descendent nucleus pulposus cells in mouse injury-induced fibrotic intervertebral discs, Aging Cell 19(11) (2020) e13254.

[26] S. Mohanty, R. Pinelli, P. Pricop, T.J. Albert, C.L. Dahia, Chondrocyte-like nested cells in the aged intervertebral disc are late-stage nucleus pulposus cells, Aging Cell 18(5) (2019) e13006.

[27] F.C. Bach, D.W. Poramba-Liyanage, F.M. Riemers, J. Guicheux, A. Camus, J.C. Iatridis, D. Chan, K. Ito, C.L. Le Maitre, M.A. Tryfonidou, Notochordal Cell-Based Treatment Strategies and Their Potential in Intervertebral Disc Regeneration, Front Cell Dev Biol 9 (2021) 780749.

[28] D.J. Aguiar, S.L. Johnson, T.R. Oegema, Notochordal cells interact with nucleus pulposus cells: regulation of proteoglycan synthesis, Exp Cell Res 246(1) (1999) 129–37.

[29] M.O. Baffi, E. Slattery, P. Sohn, H.L. Moses, A. Chytil, R. Serra, Conditional deletion of the TGF-beta type II receptor in Col2a expressing cells results in defects in the axial skeleton without alterations in chondrocyte differentiation or embryonic development of long bones, Dev Biol 276(1) (2004) 124–42.

[30] S. Chen, S. Liu, K. Ma, L. Zhao, H. Lin, Z. Shao, TGF-beta signaling in intervertebral disc health and disease, Osteoarthritis and cartilage 27(8) (2019) 1109–1117.

[31] L.J. Smith, J.A. Chiaro, N.L. Nerurkar, D.H. Cortes, S.D. Horava, N.M. Hebela, R.L. Mauck, G.R. Dodge, D.M. Elliott, Nucleus pulposus cells synthesize a functional extracellular matrix and respond to inflammatory cytokine challenge following long-term agarose culture, Eur Cell Mater 22 (2011) 291–301.

[32] K. Sun, J. Luo, J. Guo, X. Yao, X. Jing, F. Guo, The PI3K/AKT/mTOR signaling pathway in osteoarthritis: a narrative review, Osteoarthritis and cartilage 28(4) (2020) 400–409.

[33] B.A. Hemmings, D.F. Restuccia, The PI3K-PKB/Akt pathway, Cold Spring Harb Perspect Biol 7(4) (2015).

[34] Q. Xiao, Y. Teng, C. Xu, W. Pan, H. Yang, J. Zhao, Q. Zhou, Role of PI3K/AKT Signaling Pathway in Nucleus Pulposus Cells, Biomed Res Int 2021 (2021) 9941253.

[35] H. Tapp, D.J. Hernandez, P.J. Neame, T.J. Koob, Pleiotrophin inhibits chondrocyte proliferation and stimulates proteoglycan synthesis in mature bovine cartilage, Matrix Biol 18(6) (1999) 543–56.

[36] C.R. Lee, D. Sakai, T. Nakai, K. Toyama, J. Mochida, M. Alini, S. Grad, A phenotypic comparison of intervertebral disc and articular cartilage cells in the rat, Eur Spine J 16(12) (2007) 2174–85.

[37] R.J. Bai, Y.S. Li, F.J. Zhang, Osteopontin, a bridge links osteoarthritis and osteoporosis, Front Endocrinol (Lausanne) 13 (2022) 1012508.

[38] D.T. Denhardt, C.A. Lopez, E.E. Rollo, S.M. Hwang, X.R. An, S.E. Walther, Osteopontin-induced modifications of cellular functions, Ann N Y Acad Sci 760 (1995) 127–42.

[39] L. Liaw, D.E. Birk, C.B. Ballas, J.S. Whitsitt, J.M. Davidson, B.L. Hogan, Altered wound healing in mice lacking a functional osteopontin gene (spp1), J Clin Invest 101(7) (1998) 1468–78.

[40] J.M. Thayer, C.M. Giachelli, P.E. Mirkes, S.M. Schwartz, Expression of osteopontin in the head process late in gastrulation in the rat, J Exp Zool 272(3) (1995) 240–4.

[41] T. Zhou, Y. Chen, Z. Liao, L. Zhang, D. Su, Z. Li, X. Yang, X. Ke, H. Liu, Y. Chen, R. Weng, H. Shen, C. Xu, Y. Wan, R. Xu, P. Su, Spatiotemporal Characterization of Human Early Intervertebral Disc Formation at Single-Cell Resolution, Adv Sci (Weinh) 10(14) (2023) e2206296.

[42] M. Kudelko, P. Chen, V. Tam, Y. Zhang, O.Y. Kong, R. Sharma, T.Y.K. Au, M.K. To, K.S.E. Cheah, W.C.W. Chan, D. Chan, PRIMUS: Comprehensive proteomics of mouse intervertebral discs that inform novel biology and relevance to human disease modelling, Matrix Biol Plus 12 (2021) 100082.

[43] V. Tam, P. Chen, A. Yee, N. Solis, T. Klein, M. Kudelko, R. Sharma, W.C. Chan, C.M. Overall, L. Haglund, P.C. Sham, K.S.E. Cheah, D. Chan, DIPPER, a spatiotemporal proteomics atlas of human intervertebral discs for exploring ageing and degeneration dynamics, Elife 9 (2020).

[44] D. Powner, P.M. Kopp, S.J. Monkley, D.R. Critchley, F. Berditchevski, Tetraspanin CD9 in cell migration, Biochem Soc Trans 39(2) (2011) 563–7.

[45] X. Jiang, M. Teng, R. Ji, D. Zhang, Z. Zhang, Y. Lv, Q. Zhang, J. Zhang, Y. Huang, CD9 regulates keratinocyte differentiation and motility by recruiting E-cadherin to the plasma membrane and activating the PI3K/Akt pathway, Biochim Biophys Acta Mol Cell Res 1867(2) (2020) 118574.

[46] C. Brosseau, L. Colas, A. Magnan, S. Brouard, CD9 Tetraspanin: A New Pathway for the Regulation of Inflammation?, Front Immunol 9 (2018) 2316.

[47] W. Yu, L. Zhong, L. Yao, Y. Wei, T. Gui, Z. Li, H. Kim, N. Holdreith, X. Jiang, W. Tong, N. Dyment, X.S. Liu, S. Yang, Y. Choi, J. Ahn, L. Qin, Bone marrow adipogenic lineage precursors promote osteoclastogenesis in bone remodeling and pathologic bone loss, J Clin Invest 131(2) (2021).

[48] M. Kim, D.R. Steinberg, J.A. Burdick, R.L. Mauck, Extracellular vesicles mediate improved functional outcomes in engineered cartilage produced from MSC/chondrocyte cocultures, Proc Natl Acad Sci U S A 116(5) (2019) 1569–1578.

[49] S.E. Gullbrand, N.R. Malhotra, T.P. Schaer, Z. Zawacki, J.T. Martin, J.R. Bendigo, A.H. Milby, G.R. Dodge, E.J. Vresilovic, D.M. Elliott, R.L. Mauck, L.J. Smith, A large animal model that recapitulates the spectrum of human intervertebral disc degeneration, Osteoarthritis and cartilage 25(1) (2017) 146–156.

[50] L. Zhong, L. Yao, R.J. Tower, Y. Wei, Z. Miao, J. Park, R. Shrestha, L. Wang, W. Yu, N. Holdreith, X. Huang, Y. Zhang, W. Tong, Y. Gong, J. Ahn, K. Susztak, N. Dyment, M. Li, F. Long, C. Chen, P. Seale, L. Qin, Single cell transcriptomics identifies a unique adipose lineage cell population that regulates bone marrow environment, Elife 9 (2020).

[51] Y. Hao, S. Hao, E. Andersen-Nissen, W.M. Mauck, 3rd, S. Zheng, A. Butler, M.J. Lee, A.J. Wilk, C. Darby, M. Zager, P. Hoffman, M. Stoeckius, E. Papalexi, E.P. Mimitou, J. Jain, A. Srivastava, T. Stuart, L.M. Fleming, B. Yeung, A.J. Rogers, J.M. McElrath, C.A. Blish, R. Gottardo, P. Smibert, R. Satija, Integrated analysis of multimodal single-cell data, Cell 184(13) (2021) 3573–3587 e29.

[52] I. Tirosh, B. Izar, S.M. Prakadan, M.H. Wadsworth, 2nd, D. Treacy, J.J. Trombetta, A. Rotem, C. Rodman, C. Lian, G. Murphy, M. Fallahi-Sichani, K. Dutton-Regester, J.R. Lin, O. Cohen, P. Shah, D. Lu, A.S. Genshaft, T.K. Hughes, C.G. Ziegler, S.W. Kazer, A. Gaillard, K.E. Kolb, A.C. Villani, C.M. Johannessen, A.Y. Andreev, E.M. Van Allen, M. Bertagnolli, P.K. Sorger, R.J. Sullivan, K.T. Flaherty, D.T. Frederick, J. Jane-Valbuena, C.H. Yoon, O. Rozenblatt-Rosen, A.K. Shalek, A. Regev, L.A. Garraway, Dissecting the multicellular ecosystem of metastatic melanoma by single-cell RNA-seq, Science 352(6282) (2016) 189–96.

[53] C. Trapnell, D. Cacchiarelli, J. Grimsby, P. Pokharel, S. Li, M. Morse, N.J. Lennon, K.J. Livak, T.S. Mikkelsen, J.L. Rinn, The dynamics and regulators of cell fate decisions are revealed by pseudotemporal ordering of single cells, Nat Biotechnol 32(4) (2014) 381–386.

[54] V. Bergen, M. Lange, S. Peidli, F.A. Wolf, F.J. Theis, Generalizing RNA velocity to transient cell states through dynamical modeling, Nat Biotechnol 38(12) (2020) 1408–1414.

[55] D. Szklarczyk, A.L. Gable, K.C. Nastou, D. Lyon, R. Kirsch, S. Pyysalo, N.T. Doncheva, M. Legeay, T. Fang, P. Bork, L.J. Jensen, C. von Mering, The STRING database in 2021: customizable protein-protein networks, and functional characterization of user-uploaded gene/measurement sets, Nucleic Acids Res 49(D1) (2021) D605–D612.

