## Supplementary Figures and Tables for "Single Cell RNA Sequencing Reveals Emergent Notochord-Derived Cell Subpopulations in the Postnatal Nucleus Pulposus"

Supplementary Figures 1 and 2

Supplementary Tables 1-3

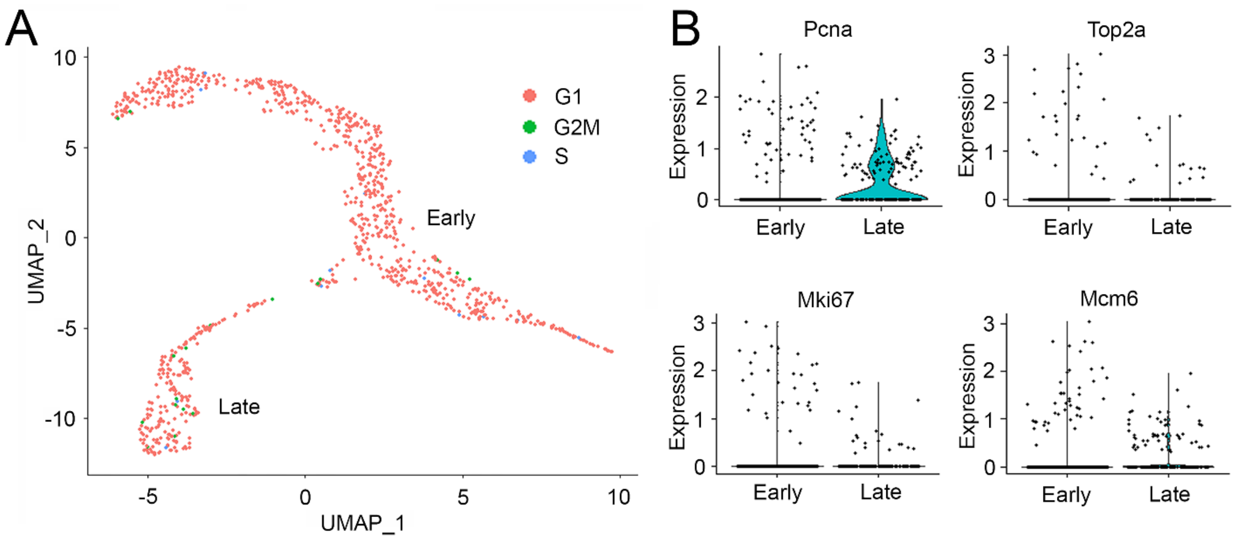


**Supplementary Figure 1**. **A.** UMAP of NP cells showing cell cycle state. **B.** Violin plots showing relative expression of cell proliferation markers in early and late stage NP cells.

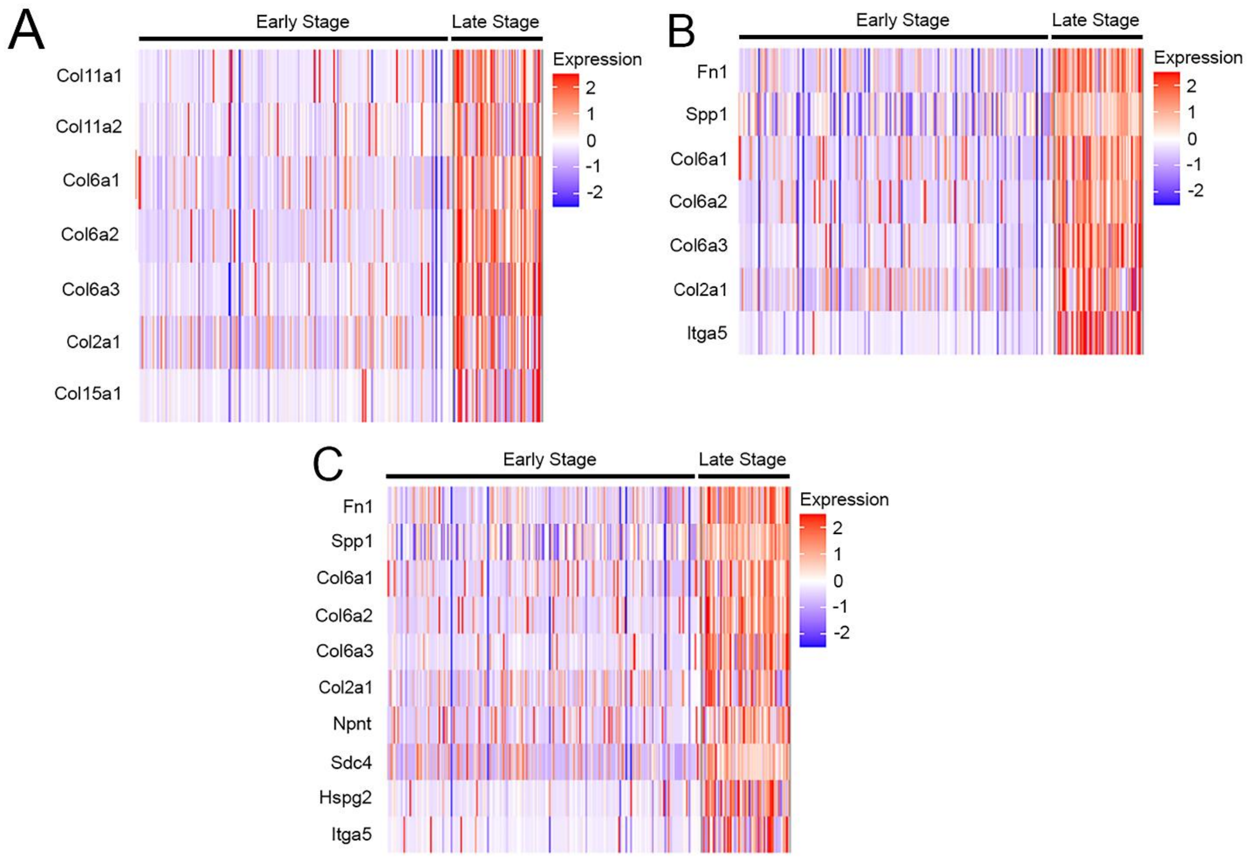


**Supplementary Figure 2.** Heatmaps showing differentially expressed genes in the **A.** Protein digestion and absorption; **B.** Focal adhesion; and **C.** ECM-receptor interactions pathways, respectively, between late and early-stage NP cells.

**Supplementary Table 1.** Genes significantly upregulated in late versus early-stage NP cells, ranked in descending order by relative log2 fold change.

| Gene | Log2 Fold Change | p (adjusted) |
|---|---|---|
| Fn1 | 4.11 | 4.26E-96 |
| Acan | 4.11 | 1.25E-100 |
| Fos | 3.32 | 2.06E-40 |
| Col2a1 | 3.25 | 2.44E-52 |
| Atf3 | 3.10 | 7.44E-57 |
| Spp1 | 2.98 | 1.89E-87 |
| Egr1 | 2.86 | 1.95E-09 |
| Fosb | 2.74 | 4.22E-61 |
| Junb | 2.69 | 2.01E-29 |
| CD109 | 2.56 | 2.21E-91 |
| Lyst | 2.53 | 5.84E-69 |
| Htra1 | 2.52 | 2.30E-89 |
| Col6a2 | 2.23 | 7.14E-83 |
| Ecrg4 | 2.23 | 5.36E-57 |
| Col6a1 | 2.20 | 6.04E-78 |
| Col11a1 | 2.16 | 3.57E-59 |
| Col6a3 | 2.10 | 1.83E-77 |
| Hapln1 | 2.10 | 1.12E-72 |
| Ccn2 | 2.09 | 2.03E-05 |
| Sparc | 2.05 | 1.52E-48 |
| Ahnak | 2.03 | 3.82E-70 |
| Mgp | 2.02 | 7.88E-38 |
| Susd5 | 2.01 | 1.58E-77 |
| Sostdc1 | 1.99 | 7.86E-43 |
| Emp1 | 1.98 | 7.24E-17 |
| Socs3 | 1.96 | 5.87E-07 |
| CD9 | 1.90 | 1.21E-35 |
| Ccn1 | 1.90 | 5.80E-24 |
| Ptgs1 | 1.89 | 1.89E-66 |
| App | 1.88 | 6.79E-76 |
| Dst | 1.87 | 1.10E-65 |
| Fmod | 1.83 | 3.00E-57 |
| Gadd45g | 1.83 | 7.51E-48 |
| Slc12a2 | 1.80 | 2.93E-53 |
| Mfge8 | 1.71 | 8.43E-50 |
| Ahnak2 | 1.69 | 1.45E-74 |
| Peg3 | 1.65 | 3.89E-50 |
| Cdkn1a | 1.65 | 1.61E-21 |
| Dusp1 | 1.63 | 2.82E-09 |

| | | |
|---|---|---|
| Cpe | 1.57 | 3.77E-60 |
| Sparcl1 | 1.56 | 5.45E-64 |
| Itm2a | 1.53 | 1.09E-58 |
| Ecm1 | 1.53 | 1.67E-48 |
| Fgl2 | 1.51 | 3.46E-33 |
| Zbtb20 | 1.49 | 2.02E-58 |
| Gm26532 | 1.47 | 9.18E-03 |
| Col11a2 | 1.46 | 6.52E-51 |
| Xylt1 | 1.45 | 4.61E-54 |
| Id1 | 1.43 | 1.27E-43 |
| Rrbp1 | 1.43 | 1.24E-67 |
| Klf4 | 1.43 | 4.97E-16 |
| Mt1 | 1.41 | 9.80E-15 |
| Ddr2 | 1.41 | 4.34E-45 |
| Cp | 1.41 | 4.57E-49 |
| Igfbp7 | 1.40 | 3.81E-55 |
| Sdc4 | 1.39 | 1.45E-49 |
| Snorc | 1.38 | 1.80E-57 |
| Id3 | 1.38 | 7.94E-19 |
| Ptn | 1.37 | 7.39E-42 |
| Meltf | 1.37 | 3.61E-28 |
| Ltbp1 | 1.37 | 2.79E-27 |
| Npnt | 1.35 | 2.94E-62 |
| Ifrd1 | 1.34 | 3.27E-14 |
| Lgmn | 1.33 | 6.40E-64 |
| Meg3 | 1.33 | 1.26E-04 |
| Cck | 1.33 | 7.61E-35 |
| Sox9 | 1.33 | 4.73E-43 |
| Kif21a | 1.32 | 7.25E-54 |
| Scd2 | 1.31 | 3.58E-51 |
| Lrp1 | 1.31 | 7.01E-45 |
| Dsp | 1.30 | 1.53E-54 |
| Ctsl | 1.28 | 1.07E-48 |
| Hspg2 | 1.27 | 1.31E-37 |
| Plod2 | 1.27 | 7.02E-44 |
| Klf9 | 1.26 | 5.81E-11 |
| Itga5 | 1.24 | 4.77E-15 |
| Cytl1 | 1.23 | 1.57E-27 |
| Ehd3 | 1.22 | 2.23E-34 |
| Zim1 | 1.21 | 9.54E-42 |
| Lmo7 | 1.20 | 8.55E-45 |
| Cspg4 | 1.18 | 3.76E-30 |
| Angptl1 | 1.17 | 7.35E-42 |
| Chrm2 | 1.16 | 1.98E-46 |
| Gas1 | 1.14 | 2.91E-39 |

| | | |
|---|---|---|
| Fat1 | 1.13 | 1.21E-30 |
| Palld | 1.12 | 5.41E-42 |
| Cdh6 | 1.12 | 4.75E-24 |
| Olfml2b | 1.11 | 1.31E-35 |
| Loxl2 | 1.11 | 5.18E-46 |
| Prnp | 1.11 | 1.41E-53 |
| Col15a1 | 1.10 | 6.28E-11 |
| Serpinh1 | 1.09 | 1.18E-47 |
| Smoc1 | 1.09 | 6.41E-49 |
| Cpm | 1.08 | 1.39E-22 |
| Csgalnact1 | 1.06 | 2.00E-49 |
| Nr4a2 | 1.05 | 3.18E-08 |
| P4ha1 | 1.05 | 1.44E-59 |
| Serpina1e | 1.05 | 5.76E-30 |
| Prelp | 1.04 | 2.22E-42 |
| Tnfrsf12a | 1.03 | 1.11E-28 |
| Ctsb | 1.03 | 1.06E-56 |
| Id2 | 1.02 | 8.34E-28 |
| Tob1 | 1.01 | 8.28E-38 |
| Pde4b | 1.00 | 7.58E-38 |
| Aebp1 | 0.99 | 2.05E-47 |
| Mxra8 | 0.97 | 4.70E-27 |
| Slc15a5 | 0.97 | 1.05E-52 |
| Sgk1 | 0.97 | 1.87E-14 |
| Tubb2a | 0.95 | 5.98E-28 |
| Timp2 | 0.95 | 3.40E-37 |
| Tnfrsf11b | 0.94 | 2.66E-21 |
| Cdh2 | 0.94 | 2.59E-46 |
| Ucma | 0.93 | 9.36E-11 |
| Plcl1 | 0.92 | 4.62E-34 |
| Tln2 | 0.92 | 6.88E-49 |
| Wwp2 | 0.91 | 3.63E-26 |
| Bhlhe40 | 0.91 | 1.03E-08 |
| Itga1 | 0.90 | 1.22E-13 |
| Cpxm2 | 0.90 | 1.73E-34 |
| Limch1 | 0.90 | 1.16E-27 |
| Slc7a2 | 0.90 | 1.49E-43 |
| Enpp2 | 0.88 | 4.95E-07 |
| Bgn | 0.88 | 3.97E-41 |
| Tgfb2 | 0.88 | 1.09E-57 |
| Pcolce2 | 0.87 | 7.08E-49 |
| Pdia6 | 0.87 | 1.63E-45 |
| Gpc3 | 0.87 | 6.55E-43 |
| Tsc22d1 | 0.86 | 2.01E-29 |
| Itih5 | 0.86 | 1.12E-08 |

| | | |
|---|---|---|
| Slc2a1 | 0.86 | 3.61E-33 |
| Tnxb | 0.85 | 1.71E-04 |
| Pkd2 | 0.84 | 5.50E-54 |
| Flt1 | 0.84 | 4.80E-22 |
| Cst3 | 0.83 | 9.18E-33 |
| Crip1 | 0.81 | 2.12E-16 |
| Fkbp9 | 0.81 | 7.17E-44 |
| Nrp2 | 0.80 | 7.26E-37 |
| Pcdh7 | 0.80 | 2.99E-24 |
| Trps1 | 0.80 | 4.45E-39 |
| Gja1 | 0.79 | 6.89E-14 |
| Pim3 | 0.78 | 1.21E-49 |
| Col9a3 | 0.77 | 2.58E-21 |
| Creb3l2 | 0.77 | 7.41E-39 |
| Hmgcs1 | 0.76 | 1.93E-18 |
| Gpx3 | 0.76 | 1.86E-07 |
| Gsn | 0.76 | 9.38E-42 |
| Fkbp10 | 0.74 | 8.76E-34 |
| Pmepa1 | 0.73 | 2.12E-16 |
| Cavin2 | 0.73 | 5.61E-33 |
| Sox6 | 0.73 | 1.31E-56 |
| Hbb-bt | 0.73 | 5.42E-04 |
| Hif1a | 0.72 | 9.24E-54 |
| Myh11 | 0.71 | 2.84E-11 |
| Tafa1 | 0.70 | 1.33E-12 |
| Lmna | 0.69 | 2.63E-24 |
| S100b | 0.69 | 8.47E-21 |
| P3h4 | 0.68 | 4.54E-27 |
| Atp6v0c | 0.68 | 6.50E-60 |
| Rnase4 | 0.68 | 8.65E-39 |
| Ptprd | 0.67 | 1.23E-18 |
| Pmp22 | 0.67 | 9.13E-39 |
| Lifr | 0.66 | 7.68E-47 |
| Hspa1a | 0.66 | 2.16E-18 |
| Sulf2 | 0.65 | 4.82E-18 |
| Plxna2 | 0.65 | 8.87E-52 |
| Cdh13 | 0.65 | 3.07E-28 |
| Fxyd3 | 0.65 | 1.84E-27 |
| Auts2 | 0.64 | 3.40E-44 |
| Cdon | 0.64 | 1.45E-21 |
| Hopx | 0.63 | 4.65E-12 |
| Cd302 | 0.63 | 3.24E-35 |
| Serpine1 | 0.62 | 2.94E-14 |
| Tm4sf1 | 0.61 | 6.15E-12 |
| Krt18 | 0.61 | 5.20E-11 |

| | | |
|---|---|---|
| Itgb5 | 0.61 | 7.36E-10 |
| S100a10 | 0.61 | 1.87E-12 |
| Ccdc80 | 0.61 | 5.99E-10 |
| Col1a1 | 0.61 | 8.39E-07 |
| Mpzl2 | 0.59 | 7.47E-08 |
| Fndc3b | 0.59 | 3.33E-25 |
| Hba-a2 | 0.58 | 1.88E-06 |
| Fermt2 | 0.58 | 2.21E-24 |
| Apoe | 0.57 | 2.44E-02 |
| Gpc6 | 0.57 | 1.60E-15 |
| Abi3bp | 0.57 | 2.10E-48 |
| Nfatc2 | 0.57 | 1.79E-48 |
| Rcn3 | 0.56 | 2.14E-25 |
| Calr | 0.56 | 2.64E-32 |
| Timp3 | 0.56 | 3.49E-32 |
| Fgfr1 | 0.55 | 1.37E-34 |
| Tmx4 | 0.54 | 7.10E-65 |
| Tgfbr3 | 0.54 | 4.15E-06 |
| Tmem255a | 0.54 | 1.95E-30 |
| Adcy5 | 0.54 | 6.11E-33 |
| Pde1a | 0.54 | 5.05E-13 |
| Pcp4l1 | 0.53 | 3.54E-23 |
| Ikbip | 0.53 | 6.06E-20 |
| Htra3 | 0.52 | 1.23E-36 |
| Crip2 | 0.52 | 4.05E-18 |
| Gpx8 | 0.51 | 8.66E-21 |
| Tspan3 | 0.51 | 3.29E-25 |
| Rgcc | 0.51 | 1.93E-02 |
| Slc26a2 | 0.51 | 2.07E-02 |
| Ctsk | 0.51 | 1.42E-06 |
| Slc20a1 | 0.50 | 1.08E-19 |
| P4ha2 | 0.49 | 8.05E-04 |
| Lypd1 | 0.48 | 3.68E-03 |
| P3h2 | 0.48 | 1.09E-35 |
| Srgn | 0.47 | 1.10E-20 |
| Gm26802 | 0.46 | 4.84E-11 |
| Itm2c | 0.46 | 6.99E-17 |
| Dnajb1 | 0.46 | 1.14E-02 |
| Btbd3 | 0.45 | 1.11E-21 |
| S100a3 | 0.45 | 2.91E-37 |
| Chst11 | 0.44 | 9.43E-30 |
| Ltbp3 | 0.44 | 9.48E-22 |
| Crtap | 0.43 | 1.05E-17 |
| Plaur | 0.43 | 3.53E-04 |
| Sema3c | 0.43 | 7.43E-15 |

| | | |
|---|---|---|
| Socs1 | 0.43 | 1.32E-34 |
| Grb10 | 0.43 | 7.72E-29 |
| Gadd45b | 0.43 | 3.05E-13 |
| Myc | 0.43 | 8.88E-08 |
| Scara3 | 0.41 | 1.34E-40 |
| Icam1 | 0.41 | 1.18E-19 |
| Fgfr3 | 0.41 | 7.53E-10 |
| Gpc1 | 0.40 | 1.08E-23 |
| Ppic | 0.39 | 4.66E-02 |
| Sgms2 | 0.39 | 3.42E-20 |
| Sbsn | 0.38 | 4.61E-07 |
| Ptdss2 | 0.38 | 6.57E-63 |
| Maged2 | 0.37 | 1.05E-13 |
| Adamts2 | 0.37 | 7.06E-06 |
| Fkbp14 | 0.37 | 1.23E-23 |
| Cav2 | 0.37 | 7.52E-19 |
| Azgp1 | 0.36 | 6.10E-22 |
| Srpx2 | 0.36 | 5.08E-07 |
| C1galt1c1 | 0.36 | 5.80E-03 |
| Mageh1 | 0.35 | 9.09E-22 |
| Clu | 0.35 | 1.17E-13 |
| Npdc1 | 0.34 | 2.29E-27 |
| Tmod2 | 0.34 | 3.06E-12 |
| Plpp1 | 0.34 | 1.26E-08 |
| Slit3 | 0.34 | 4.42E-13 |
| Nfix | 0.34 | 9.53E-19 |
| Tmem176b | 0.33 | 2.04E-15 |
| Lamb2 | 0.33 | 1.91E-04 |
| Bhlhe41 | 0.32 | 6.47E-03 |
| Sobp | 0.32 | 4.57E-39 |
| Ptgds | 0.32 | 5.71E-44 |
| Raph1 | 0.31 | 9.85E-30 |
| Kdelr3 | 0.31 | 1.76E-03 |
| Selenbp1 | 0.30 | 7.05E-13 |
| Sfrp5 | 0.30 | 1.24E-22 |
| Ccn5 | 0.29 | 4.60E-04 |
| Maf | 0.29 | 7.09E-11 |
| Ifi27l2a | 0.29 | 3.81E-02 |
| Id4 | 0.29 | 1.11E-02 |
| 9330158H04Rik | 0.29 | 3.95E-21 |
| Lmcd1 | 0.29 | 8.36E-43 |
| Aspa | 0.28 | 1.15E-27 |
| Fgfr2 | 0.28 | 2.60E-35 |
| Pak3 | 0.28 | 3.72E-45 |
| Itpkc | 0.28 | 1.20E-29 |

| | | |
|---|---|---|
| Gfod2 | 0.28 | 2.10E-75 |
| Rnf149 | 0.27 | 5.74E-55 |
| Thbs3 | 0.26 | 1.63E-07 |
| Rhoj | 0.26 | 3.49E-10 |
| Efemp2 | 0.26 | 1.68E-23 |
| Bmp1 | 0.25 | 3.58E-56 |
| Cxcl2 | 0.25 | 1.18E-16 |
| Irx5 | 0.25 | 2.76E-06 |
| Slurp1 | 0.25 | 1.86E-31 |
| Egr4 | 0.25 | 2.48E-35 |

**Supplementary Table 2.** Genes significantly downregulated in late versus early-stage NP cells, ranked in descending order by relative log2 fold change.

| Gene | Log2 Fold Change | p (adjusted) |
|---|---|---|
| Rbp4 | -1.63 | 2.50E-18 |
| Cdo1 | -1.35 | 5.85E-07 |
| Plp1 | -1.18 | 2.27E-37 |
| Cox8b | -0.99 | 6.77E-61 |
| 3110079O15Rik | -0.92 | 2.41E-02 |
| Hagh | -0.80 | 1.00E-04 |
| Mif | -0.70 | 8.47E-18 |
| Npm1 | -0.70 | 1.96E-04 |
| Mt3 | -0.69 | 2.06E-66 |
| 1500011K16Rik | -0.69 | 2.54E-04 |
| Glrx5 | -0.63 | 2.00E-15 |
| Rbp1 | -0.62 | 1.09E-13 |
| Car3 | -0.61 | 6.69E-12 |
| Plac8 | -0.61 | 5.76E-08 |
| Myl9 | -0.57 | 7.17E-22 |
| Fabp5 | -0.56 | 2.90E-18 |
| Blvrb | -0.55 | 1.27E-06 |
| 1500015O10Rik | -0.55 | 3.40E-13 |
| Nhp2 | -0.53 | 1.17E-06 |
| Fxyd6 | -0.51 | 3.74E-03 |
| Hebp1 | -0.51 | 1.62E-08 |
| Car2 | -0.49 | 8.35E-11 |
| Sln | -0.42 | 6.51E-08 |
| Vat1 | -0.40 | 6.75E-03 |
| Car1 | -0.40 | 1.39E-52 |
| 1110008F13Rik | -0.36 | 9.73E-07 |
| Fkbp4 | -0.36 | 3.75E-04 |
| Col22a1 | -0.36 | 2.71E-20 |
| Ndufa4l2 | -0.35 | 8.70E-03 |
| Nkg7 | -0.35 | 9.85E-60 |
| Ncl | -0.32 | 9.08E-04 |
| Wfdc21 | -0.31 | 5.92E-12 |
| Timm8a1 | -0.30 | 5.03E-09 |
| Pla2g12a | -0.29 | 3.11E-07 |
| Ctse | -0.28 | 8.89E-70 |
| Mustn1 | -0.28 | 1.53E-21 |
| Apex1 | -0.27 | 1.13E-03 |
| Isg20 | -0.27 | 6.93E-03 |
| Clec12a | -0.27 | 1.98E-29 |

| | | |
|---|---|---|
| Ifrd2 | -0.26 | 1.25E-46 |
| Hist1h4i | -0.26 | 3.27E-07 |
| Col1a2 | -0.25 | 5.02E-08 |
| Stard10 | -0.25 | 3.31E-48 |

**Supplementary Table 3**. Cell-cell interactions identified using CellChat.

| Source | Target | Annotation | Ligand | Receptor | Probability | Interaction Name | Evidence |
|---|---|---|---|---|---|---|---|
| Early stage NP cells | Late stage NP cells | Secreted Signaling | Ptn | Sdc2 | 0.007 | Ptn - Sdc2 | PMID: 28356350; PMID: 25620911 |
| Early stage NP cells | Late stage NP cells | Secreted Signaling | Ptn | Sdc4 | 0.096 | Ptn - Sdc4 | PMID: 28356350; PMID: 25620911 |
| Early stage NP cells | Late stage NP cells | Secreted Signaling | Ptn | Ncl | 0.051 | Ptn - Ncl | PMID: 28356350; PMID: 25620911 |
| Early stage NP cells | Late stage NP cells | Secreted Signaling | Spp1 | Cd44 | 0.011 | Spp1 - Cd44 | PMID: 21907263 |
| Early stage NP cells | Late stage NP cells | Secreted Signaling | Spp1 | Itgav+Itgb1 | 0.054 | Spp1 - (Itgav+Itgb1) | PMID: 21907263 |
| Early stage NP cells | Late stage NP cells | Secreted Signaling | Spp1 | Itgav+Itgb5 | 0.028 | Spp1 - (Itgav+Itgb5) | PMID: 21907263 |
| Early stage NP cells | Late stage NP cells | Secreted Signaling | Spp1 | Itga5+Itgb1 | 0.092 | Spp1 - (Itga5+Itgb1) | PMID: 21907263 |
| Late stage NP cells | Early stage NP cells | ECM-Receptor | Col1a1 | Sdc4 | 0.009 | Col1a1 - Sdc4 | KEGG: mmu04512 |
| Late stage NP cells | Early stage NP cells | ECM-Receptor | Col1a2 | Sdc4 | 0.002 | Col1a2 - Sdc4 | KEGG: mmu04512 |
| Late stage NP cells | Early stage NP cells | ECM-Receptor | Col2a1 | Sdc4 | 0.026 | Col2a1 - Sdc4 | KEGG: mmu04512 |
| Late stage NP cells | Early stage NP cells | ECM-Receptor | Col6a1 | Sdc4 | 0.028 | Col6a1 - Sdc4 | KEGG: mmu04512 |
| Late stage NP cells | Early stage NP cells | ECM-Receptor | Col6a2 | Sdc4 | 0.025 | Col6a2 - Sdc4 | KEGG: mmu04512 |
| Late stage NP cells | Early stage NP cells | ECM-Receptor | Col6a3 | Sdc4 | 0.02 | Col6a3 - Sdc4 | KEGG: mmu04512 |
| Late stage NP cells | Early stage NP cells | ECM-Receptor | Col9a3 | Sdc4 | 0.003 | Col9a3 - Sdc4 | KEGG: mmu04512 |
| Late stage NP cells | Early stage NP cells | ECM-Receptor | Fn1 | Sdc4 | 0.048 | Fn1 - Sdc4 | KEGG: mmu04512 |

| | | | | | | | |
|---|---|---|---|---|---|---|---|
| Late stage NP cells | Early stage NP cells | ECM-Receptor | Thbs3 | Sdc4 | 0.001 | Thbs3 - Sdc4 | KEGG: mmu04512 |
| Late stage NP cells | Early stage NP cells | ECM-Receptor | Tnxb | Sdc4 | 0.008 | Tnxb - Sdc4 | KEGG: mmu04512 |
| Late stage NP cells | Early stage NP cells | Secreted Signaling | Ptn | Sdc4 | 0.037 | Ptn - Sdc4 | PMID: 28356350; PMID: 25620911 |
| Late stage NP cells | Early stage NP cells | Secreted Signaling | Ptn | Ncl | 0.035 | Ptn - Ncl | PMID: 28356350; PMID: 25620911 |
| Late stage NP cells | Late stage NP cells | Cell-Cell Contact | Cdh2 | Cdh2 | 0.048 | Cdh2 - Cdh2 | KEGG: mmu04514 |
| Late stage NP cells | Late stage NP cells | Cell-Cell Contact | Dsg2 | Dsc3 | 0.001 | Dsg2 - Dsc3 | PMID: 27298358 |
| Late stage NP cells | Late stage NP cells | Cell-Cell Contact | Ncam1 | Fgfr1 | 0.004 | Ncam1 - Fgfr1 | PMID: 12791257 |
| Late stage NP cells | Late stage NP cells | Cell-Cell Contact | Ncam1 | Ncam1 | 0.013 | Ncam1 - Ncam1 | KEGG: mmu04514 |
| Late stage NP cells | Late stage NP cells | ECM-Receptor | Col1a1 | Itga1+Itgb1 | 0.025 | Col1a1 - (Itga1+Itgb1) | KEGG: mmu04512 |
| Late stage NP cells | Late stage NP cells | ECM-Receptor | Col1a1 | Itga10+Itgb1 | 0.005 | Col1a1 - (Itga10+Itgb1) | KEGG: mmu04512 |
| Late stage NP cells | Late stage NP cells | ECM-Receptor | Col1a1 | Cd44 | 0.003 | Col1a1 - Cd44 | KEGG: mmu04512 |
| Late stage NP cells | Late stage NP cells | ECM-Receptor | Col1a1 | Sdc4 | 0.055 | Col1a1 - Sdc4 | KEGG: mmu04512 |
| Late stage NP cells | Late stage NP cells | ECM-Receptor | Col1a2 | Itga1+Itgb1 | 0.005 | Col1a2 - (Itga1+Itgb1) | KEGG: mmu04512 |
| Late stage NP cells | Late stage NP cells | ECM-Receptor | Col1a2 | Itga10+Itgb1 | 0.001 | Col1a2 - (Itga10+Itgb1) | KEGG: mmu04512 |
| Late stage NP cells | Late stage NP cells | ECM-Receptor | Col1a2 | Cd44 | 0.001 | Col1a2 - Cd44 | KEGG: mmu04512 |
| Late stage NP cells | Late stage NP cells | ECM-Receptor | Col1a2 | Sdc4 | 0.012 | Col1a2 - Sdc4 | KEGG: mmu04512 |

| | | | | | | | |
|---|---|---|---|---|---|---|---|
| Late stage NP cells | Late stage NP cells | ECM-Receptor | Col2a1 | Itga1+Itgb1 | 0.068 | Col2a1 - (Itga1+Itgb1) | KEGG: mmu04512 |
| Late stage NP cells | Late stage NP cells | ECM-Receptor | Col2a1 | Itga10+Itgb1 | 0.013 | Col2a1 - (Itga10+Itgb1) | KEGG: mmu04512 |
| Late stage NP cells | Late stage NP cells | ECM-Receptor | Col2a1 | Cd44 | 0.008 | Col2a1 - Cd44 | KEGG: mmu04512 |
| Late stage NP cells | Late stage NP cells | ECM-Receptor | Col2a1 | Sdc4 | 0.143 | Col2a1 - Sdc4 | KEGG: mmu04512 |
| Late stage NP cells | Late stage NP cells | ECM-Receptor | Col6a1 | Itga1+Itgb1 | 0.074 | Col6a1 - (Itga1+Itgb1) | KEGG: mmu04512 |
| Late stage NP cells | Late stage NP cells | ECM-Receptor | Col6a1 | Itga10+Itgb1 | 0.014 | Col6a1 - (Itga10+Itgb1) | KEGG: mmu04512 |
| Late stage NP cells | Late stage NP cells | ECM-Receptor | Col6a1 | Cd44 | 0.008 | Col6a1 - Cd44 | KEGG: mmu04512 |
| Late stage NP cells | Late stage NP cells | ECM-Receptor | Col6a1 | Sdc4 | 0.154 | Col6a1 - Sdc4 | KEGG: mmu04512 |
| Late stage NP cells | Late stage NP cells | ECM-Receptor | Col6a2 | Itga1+Itgb1 | 0.066 | Col6a2 - (Itga1+Itgb1) | KEGG: mmu04512 |
| Late stage NP cells | Late stage NP cells | ECM-Receptor | Col6a2 | Itga10+Itgb1 | 0.013 | Col6a2 - (Itga10+Itgb1) | KEGG: mmu04512 |
| Late stage NP cells | Late stage NP cells | ECM-Receptor | Col6a2 | Cd44 | 0.007 | Col6a2 - Cd44 | KEGG: mmu04512 |
| Late stage NP cells | Late stage NP cells | ECM-Receptor | Col6a2 | Sdc4 | 0.14 | Col6a2 - Sdc4 | KEGG: mmu04512 |
| Late stage NP cells | Late stage NP cells | ECM-Receptor | Col6a3 | Itga1+Itgb1 | 0.053 | Col6a3 - (Itga1+Itgb1) | KEGG: mmu04512 |
| Late stage NP cells | Late stage NP cells | ECM-Receptor | Col6a3 | Itga10+Itgb1 | 0.01 | Col6a3 - (Itga10+Itgb1) | KEGG: mmu04512 |
| Late stage NP cells | Late stage NP cells | ECM-Receptor | Col6a3 | Cd44 | 0.006 | Col6a3 - Cd44 | KEGG: mmu04512 |
| Late stage NP cells | Late stage NP cells | ECM-Receptor | Col6a3 | Sdc4 | 0.113 | Col6a3 - Sdc4 | KEGG: mmu04512 |

| | | | | | | | |
|---|---|---|---|---|---|---|---|
| Late stage NP cells | Late stage NP cells | ECM-Receptor | Col9a3 | Itga1+Itgb1 | 0.009 | Col9a3 - (Itga1+Itgb1) | KEGG: mmu04512 |
| Late stage NP cells | Late stage NP cells | ECM-Receptor | Col9a3 | Itga10+Itgb1 | 0.002 | Col9a3 - (Itga10+Itgb1) | KEGG: mmu04512 |
| Late stage NP cells | Late stage NP cells | ECM-Receptor | Col9a3 | Cd44 | 0.001 | Col9a3 - Cd44 | KEGG: mmu04512 |
| Late stage NP cells | Late stage NP cells | ECM-Receptor | Col9a3 | Sdc4 | 0.02 | Col9a3 - Sdc4 | KEGG: mmu04512 |
| Late stage NP cells | Late stage NP cells | ECM-Receptor | Fn1 | Itga5+Itgb1 | 0.12 | Fn1 - (Itga5+Itgb1) | KEGG: mmu04512 |
| Late stage NP cells | Late stage NP cells | ECM-Receptor | Fn1 | Itgav+Itgb1 | 0.072 | Fn1 - (Itgav+Itgb1) | KEGG: mmu04512 |
| Late stage NP cells | Late stage NP cells | ECM-Receptor | Fn1 | Cd44 | 0.014 | Fn1 - Cd44 | KEGG: mmu04512 |
| Late stage NP cells | Late stage NP cells | ECM-Receptor | Fn1 | Sdc4 | 0.239 | Fn1 - Sdc4 | KEGG: mmu04512 |
| Late stage NP cells | Late stage NP cells | ECM-Receptor | Hspg2 | Dag1 | 0.019 | Hspg2 - Dag1 | KEGG: mmu04512 |
| Late stage NP cells | Late stage NP cells | ECM-Receptor | Lama4 | Itga1+Itgb1 | 0.005 | Lama4 - (Itga1+Itgb1) | KEGG: mmu04512 |
| Late stage NP cells | Late stage NP cells | ECM-Receptor | Lama4 | Cd44 | 0.001 | Lama4 - Cd44 | KEGG: mmu04512 |
| Late stage NP cells | Late stage NP cells | ECM-Receptor | Lama4 | Dag1 | 0.003 | Lama4 - Dag1 | KEGG: mmu04512 |
| Late stage NP cells | Late stage NP cells | ECM-Receptor | Lamb2 | Itga1+Itgb1 | 0.004 | Lamb2 - (Itga1+Itgb1) | KEGG: mmu04512 |
| Late stage NP cells | Late stage NP cells | ECM-Receptor | Lamb2 | Cd44 | 0 | Lamb2 - Cd44 | KEGG: mmu04512 |
| Late stage NP cells | Late stage NP cells | ECM-Receptor | Lamb2 | Dag1 | 0.002 | Lamb2 - Dag1 | KEGG: mmu04512 |
| Late stage NP cells | Late stage NP cells | ECM-Receptor | Lamb3 | Itga1+Itgb1 | 0.02 | Lamb3 - (Itga1+Itgb1) | KEGG: mmu04512 |

| | | | | | | | |
|---|---|---|---|---|---|---|---|
| Late stage NP cells | Late stage NP cells | ECM-Receptor | Lamb3 | Cd44 | 0.002 | Lamb3 - Cd44 | KEGG: mmu04512 |
| Late stage NP cells | Late stage NP cells | ECM-Receptor | Lamb3 | Dag1 | 0.011 | Lamb3 - Dag1 | KEGG: mmu04512 |
| Late stage NP cells | Late stage NP cells | ECM-Receptor | Lamc1 | Itga1+Itgb1 | 0.005 | Lamc1 - (Itga1+Itgb1) | KEGG: mmu04512 |
| Late stage NP cells | Late stage NP cells | ECM-Receptor | Lamc1 | Cd44 | 0.001 | Lamc1 - Cd44 | KEGG: mmu04512 |
| Late stage NP cells | Late stage NP cells | ECM-Receptor | Lamc1 | Dag1 | 0.003 | Lamc1 - Dag1 | KEGG: mmu04512 |
| Late stage NP cells | Late stage NP cells | ECM-Receptor | Thbs3 | Sdc4 | 0.009 | Thbs3 - Sdc4 | KEGG: mmu04512 |
| Late stage NP cells | Late stage NP cells | ECM-Receptor | Thbs3 | Cd47 | 0.003 | Thbs3 - Cd47 | KEGG: mmu04512 |
| Late stage NP cells | Late stage NP cells | ECM-Receptor | Tnxb | Sdc4 | 0.048 | Tnxb - Sdc4 | KEGG: mmu04512 |
| Late stage NP cells | Late stage NP cells | Secreted Signaling | Angptl1 | Itga1+Itgb1 | 0.056 | Angptl1 - (Itga1+Itgb1) | PMID: 24478758 |
| Late stage NP cells | Late stage NP cells | Secreted Signaling | Angptl2 | Itga5+Itgb1 | 0.004 | Angptl2 - (Itga5+Itgb1) | PMID: 24478758 |
| Late stage NP cells | Late stage NP cells | Secreted Signaling | Bmp5 | Bmpr1a+Acvr2a | 0.001 | Bmp5 - (Bmpr1a+Acvr2a) | KEGG: mmu04350; PMID:26893264 |
| Late stage NP cells | Late stage NP cells | Secreted Signaling | Bmp5 | Bmpr1a+Bmpr2 | 0 | Bmp5 - (Bmpr1a+Bmpr2) | KEGG: mmu04350; PMID:26893264 |
| Late stage NP cells | Late stage NP cells | Secreted Signaling | Fgf1 | Fgfr1 | 0.003 | Fgf1 - Fgfr1 | PMC: 4393358 |
| Late stage NP cells | Late stage NP cells | Secreted Signaling | Fgf1 | Fgfr2 | 0.002 | Fgf1 - Fgfr2 | PMC: 4393358 |
| Late stage NP cells | Late stage NP cells | Secreted Signaling | Fgf1 | Fgfr3 | 0.003 | Fgf1 - Fgfr3 | PMC: 4393358 |

| | | | | | | | |
|---|---|---|---|---|---|---|---|
| Late stage NP cells | Late stage NP cells | Secreted Signaling | Grn | Sort1 | 0.002 | Grn - Sort1 | PMID: 29555433 |
| Late stage NP cells | Late stage NP cells | Secreted Signaling | Hbegf | Egfr | 0.001 | Hbegf - Egfr | KEGG: mmu04012 |
| Late stage NP cells | Late stage NP cells | Secreted Signaling | Nampt | Itga5+Itgb1 | 0.003 | Nampt - (Itga5+Itgb1) | PMID: 28490838 |
| Late stage NP cells | Late stage NP cells | Secreted Signaling | Ptn | Sdc2 | 0.015 | Ptn - Sdc2 | PMID: 28356350; PMID: 25620911 |
| Late stage NP cells | Late stage NP cells | Secreted Signaling | Ptn | Sdc4 | 0.193 | Ptn - Sdc4 | PMID: 28356350; PMID: 25620911 |
| Late stage NP cells | Late stage NP cells | Secreted Signaling | Ptn | Ncl | 0.108 | Ptn - Ncl | PMID: 28356350; PMID: 25620911 |
| Late stage NP cells | Late stage NP cells | Secreted Signaling | Sema3b | Nrp2+Plxna2 | 0.002 | Sema3b - (Nrp2+Plxna2) | PMID: 27533782 |
| Late stage NP cells | Late stage NP cells | Secreted Signaling | Sema3c | Nrp2+Plxna2 | 0.002 | Sema3c - (Nrp2+Plxna2) | PMID: 27533782 |
| Late stage NP cells | Late stage NP cells | Secreted Signaling | Spp1 | Cd44 | 0.018 | Spp1 - Cd44 | PMID: 21907263 |
| Late stage NP cells | Late stage NP cells | Secreted Signaling | Spp1 | Itgav+Itgb1 | 0.09 | Spp1 - (Itgav+Itgb1) | PMID: 21907263 |
| Late stage NP cells | Late stage NP cells | Secreted Signaling | Spp1 | Itgav+Itgb5 | 0.047 | Spp1 - (Itgav+Itgb5) | PMID: 21907263 |
| Late stage NP cells | Late stage NP cells | Secreted Signaling | Spp1 | Itga5+Itgb1 | 0.149 | Spp1 - (Itga5+Itgb1) | PMID: 21907263 |
| Late stage NP cells | Late stage NP cells | Secreted Signaling | Tnfsf12 | Tnfrsf12a | 0.003 | Tnfsf12 - Tnfrsf12a | KEGG: mmu04060 |
| Late stage NP cells | Late stage NP cells | Secreted Signaling | Vegfa | Flt1 | 0.004 | Vegfa - Vegfr1 | KEGG: mmu04370; PMID: 16633338 |
| Late stage NP cells | Late stage NP cells | Secreted Signaling | Vegfb | Flt1 | 0.001 | Vegfb - Vegfr1 | KEGG: mmu04370; PMID: 16633338 |
